# Toxoplasma ERK7 defends the apical complex from premature degradation

**DOI:** 10.1101/2021.12.09.471932

**Authors:** William J. O’Shaughnessy, Xiaoyu Hu, Sarah Ana Henriquez, Michael L. Reese

**Affiliations:** Department of Pharmacology, University of Texas, Southwestern Medical Center, Dallas, TX USA; Department of Biochemistry, University of Texas, Southwestern Medical Center, Dallas, TX USA

**Keywords:** Kinase, cilium, apical complex, microtubule, ubiquitin-mediated degradation

## Abstract

Accurate cellular replication balances the biogenesis and turnover of complex structures. In the apicomplexan parasite *Toxoplasma gondii,* daughter cells form within an intact mother cell, creating additional challenges to ensuring fidelity of division. The apical complex is critical to parasite infectivity and consists of apical secretory organelles and specialized cytoskeletal structures. We previously identified the kinase ERK7 as required for maturation of the apical complex in *Toxoplasma*. Here we define the *Toxoplasma* ERK7 interactome, including a putative E3 ligase, CSAR1. Genetic disruption of CSAR1 fully suppresses loss of the apical complex upon ERK7 knockdown. Furthermore, we show that CSAR1 is normally responsible for turnover of maternal cytoskeleton during cytokinesis, and that its aberrant function is driven by mislocalization from the parasite residual body to the apical complex. These data identify a protein homeostasis pathway critical for *Toxoplasma* replication and fitness and suggest an unappreciated role for the parasite residual body in compartmentalizing processes that threaten the fidelity of parasite development.

## Introduction

Cellular division is the fundamental biological process of replicating both the genetic material and the cellular structures required for viability. In most cells, successful division requires a careful balance of the creation and destruction of specific structures, and both biogenesis and turnover are highly regulated processes. Apicomplexan parasites, which include the causative agents of malaria, cryptosporidiosis, and toxoplasmosis, use a variety of extraordinary replication paradigms in which a varied number of daughter cells are fully formed within a mother parasite (1–3). In these organisms, organellar biogenesis is tightly temporally coupled to the cell cycle to ensure correct packaging into daughters (4, 5). In addition, maternal material, such as cytoskeleton, must be turned over to make room for the daughter cells’ components (6–9).

Asexual division in the parasite *Toxoplasma gondii* occurs through endodyogeny, in which two daughter cells fully form within a mother cell (Figure 1A). By maintaining an intact mother cell until cytokinesis, endodyogeny allows the parasite to remain invasive until the final steps in division. During cytokinesis, discarded maternal organelles and cytoplasm are packaged into a poorly understood organelle called the residual body (10–12, 4) (Figure 1A), through which individual parasites remain connected and share cytosolic material (13, 14). While major inroads have been made in our understanding of centrosome division (15, 16), organellar replication (5, 14, 17, 18), and cell cycle control (19–21), the process of cytokinesis has been largely unstudied in Apicomplexa.

**Figure 1:**
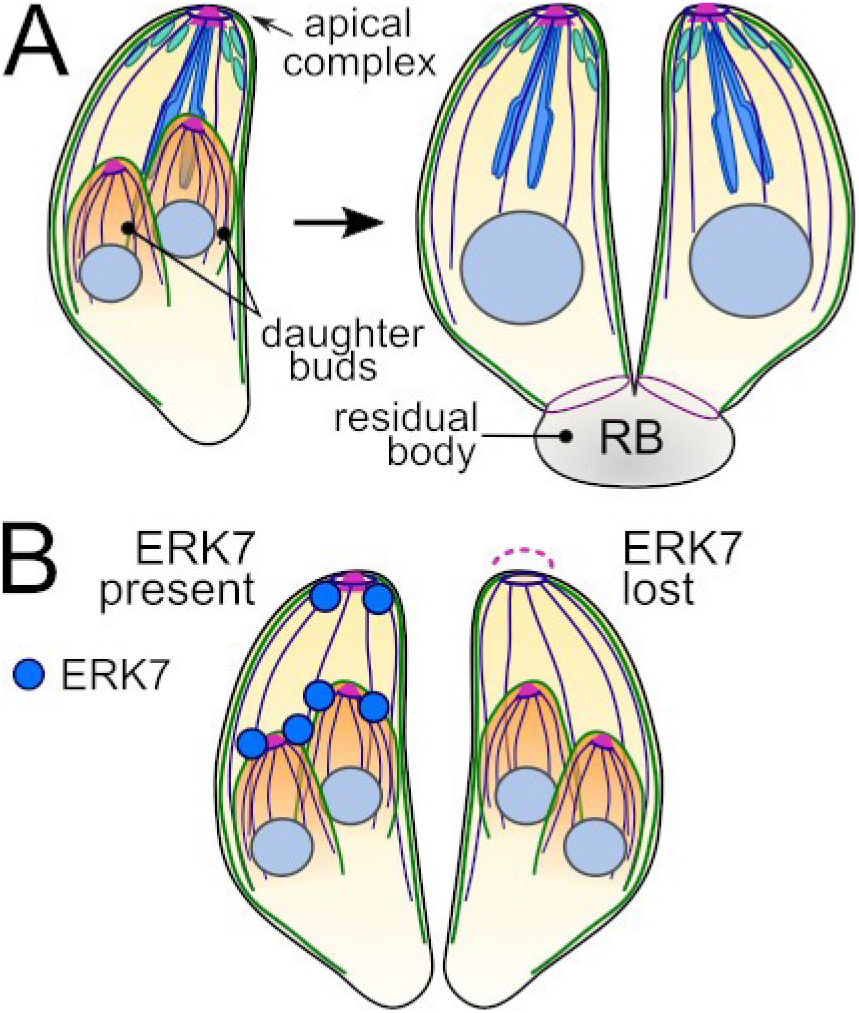
Toxoplasma divides by endodyogeny. (A) In normal Toxoplasma division, two daughter parasites are formed within the intact mother cell and are delineated a specialized cytoskeleton and membrane structure called the inner membrane complex (IMC; green). During cytokinesis, the daughter cells integrate their cytoskeleton into the mother plasma membrane. After division, the parasites remain connected through the “residual body.” (B) Diagram of ERK7 kinase loss-of-function phenotype in Toxoplasma division. When parasites divide without functional ERK7 kinase localized at their apical tips, the conoid (magenta) is lost in the mature parasite after cytokinesis, rendering the parasites noninvasive. Note that new daughter parasites grow and develop with a visible conoid.

Central to apicomplexan parasitism is a cytoskeletal structure called the apical complex, for which this phylum of parasites was named. This structure organizes specialized secretory organelles and forms the core of the parasite invasion machinery. The organizing core of the apical complex is called the conoid. Some apicomplexans, such as *Plasmodium* and *Babesia,* have lost many components of the conoid, which led to the idea that the structure is missing in these species. However, the conoid is now understood to be conserved throughout Apicomplexa (22, 23), as well as outside of Apicomplexa in early-branching Alveolates (24, 25), suggesting a truly ancient origin. In *Toxoplasma*, the conoid is formed from unusual tubulin fibers that do not form closed tubes (26) and is thought to have evolved from a more typical eukaryotic cilium, as it contains orthologs of cilium-associated proteins (27–31). The conoid is the structure through which secretion is thought to occur (32, 33), and appears essential for the initiation of parasite actin-based motility and host cell invasion (14, 34).

A scaffold of subpellicular microtubules defines the shape of both the *Toxoplasma* mother cell and daughter buds (Figure 1A). These microtubules extend from the apical tip of the daughter buds and lengthen as the daughter parasites mature. Unlike typical microtubules, the mature subpellicular microtubules are not dynamic, and are resistant to treatments that cause most other microtubule structures to disassemble, such as microtubule-depolymerizing agents, cold, and detergent-extraction (35, 36). During division, both the apical complex cytoskeleton and the subpellicular microtubules must be removed from the mother cell membrane to make room for the daughter material. How this occurs is an open question.

We recently identified the MAP kinase ERK7 as required for the development of a functional conoid in *Toxoplasma (37)*. Parasites in which ERK7 is inducibly degraded using an auxin-inducible degron (AID) (38) lose their conoids late in daughter cell assembly (Figure 1B). ERK7 localizes to the apical tip of both mature and developing daughter parasites, and loss of scaffolds required for this apical targeting phenocopies the degradation of ERK7 (39, 40). Furthermore, ERK7 kinase activity is required for its essential functions (37). Together, these data led us to the hypothesis that ERK7 phosphorylates one or more proteins that are essential to the assembly and/or maturation of the developing conoid (Figure 1B).

ERK7 is the earliest branching MAP kinase in eukaryotes, and is found in all organisms with ciliated cells (37, 41). However, very few ERK7 interacting partners or substrates have been identified and validated in any organism. In this study, we combined proximity biotinylation (42) and yeast-two-hybrid (43) to describe the *Toxoplasma* ERK7 interactome. Among the interacting proteins, we identified a number of candidate ERK7 partners with potential roles in the parasite cytoskeleton and in intracellular trafficking. Intriguingly, we identified a parasite-specific predicted RING-family ubiquitin E3 ligase that mislocalizes to the apical cap and conoid upon ERK7 degradation. We generated knockouts for the E3 ligase, and found that these parasites were unable to turnover their maternal apical complex and microtubule cytoskeleton. We therefore named the protein CSAR1 (<u>C</u>ytoskeleton <u>Sa</u>lvaging <u>R</u>ING family member 1). Most importantly, knockout of CSAR1 suppressed the phenotype of loss of ERK7 function, allowing the maturation of functional conoids even when ERK7 had been degraded. These findings demonstrate that protein degradation is a critical component of the functional maturation of the apical complex, and that ERK7 is an essential regulator of this process.

## Results

### BioID and Yeast-two-Hybrid reveal overlapping, but distinct, sets of interacting partners for Toxoplasma ERK7

The canonical MAPK pathways in fungal and animal model organisms were originally elucidated by a combination of genetic screens and interactome studies. We had previously identified apical cap protein 9 (AC9) as an essential regulatory scaffold that was required to recruit ERK7 to the parasite apical cap and thus for the formation of a functional apical complex (39). Moreover, we had demonstrated that ERK7 kinase activity is required for its function (37). Taken together, these data suggested that delineating the ERK7 interactome would identify other upstream regulators and the downstream substrates through which ERK7 mediates its function in ensuring maturation of the *Toxoplasma* apical complex. Because the conoid disappears upon auxin-induced ERK7 degradation (37), we reasoned that global phosphoproteomic analysis would be complicated by potential artifacts and secondary effects, again highlighting the need for an ERK7 interactome. To this end, we combined two orthogonal methods to identify *Toxoplasm*a proteins that interact with ERK7: yeast-two-hybrid (43) and proximity biotinylation (44). Proximity biotinylation, or BioID, uses a promiscuous biotin ligase to biotinylate proteins within ~10 nm of the bait (45), and is thus able to report on direct and indirect interactions within larger complexes as well as transient interactions, such as enzyme:substrate pairs. Yeast-two-hybrid relies on heterologous expression of a cDNA library in *S. cerevisiae* and probes only proteins that directly interact with the bait.

To perform BioID, we used a CRISPR-assisted homologous recombination strategy to C-terminally tag the endogenous copy of ERK7 with a 3xHA-BioID2 (46). We also created a control strain in which the ERK7 promoter drives the expression of mVenus-3xHA-BioID2. We infected human foreskin fibroblasts (HFF) with each of these strains, and grew them for 36 h in the presence of 150 μm biotin. Parasites were collected, lysed in RIPA buffer, and biotinylated proteins enriched by incubation with streptavidin resin. Proteins were eluted from the resin and analyzed by liquid chromatography-tandem mass spectrometry. Data from 3 replicates of each the ERK7 and control samples were compared and proteins with an average >2-fold enrichment over the control were considered candidate interactors (Table 1, Supplemental Data S1; Figure S1). Candidates with cell-cycle dependence of transcript levels (47) that showed high correlation with ERK7 or with known components of the conoid were prioritized. We also included candidates identified in our data set that had been previously identified in a previous conoid proteome (48), but for which no localization data were available. Our BioID data appeared of high quality, as the top candidates included many known components of the apical cap and IMC, including both AC9 and AC10, which we have previously identified as scaffolds that recruit ERK7 to the apical cap (39, 49).

**Table 1.** Top hits from the ERK7 interactome.

| **Known localization** |  |  |  |  |
| --- | --- | --- | --- | --- |
| **Y2H** | **BioID** | **Gene ID** | **Gene Name** | **Localization** |
| ✓ | ✓ | TG*_246950 | AC9 | Apical cap (39, 40, 69) |
| ✓ | ✓ | TG*_292950 | AC10 | Apical cap (40, 49) |
|  | ✓ | TG*_250820 | AC2 | Apical cap (64) |
|  | ✓ | TG*_214880 | AC4 | Apical cap (64) |
|  | ✓ | TG*_229640 | AC8 | Apical cap (69) |
|  | ✓ | TG*_310030 | TgCAP | Apical cap (65, 70) |
|  | ✓ | TG*_310930 | TgFBXO1 | IMC (71) |
|  | ✓ | TG*_316540 | ISP3 | IMC (64) |
|  | ✓ | TG*_231640 | IMC1 | IMC (72, 73) |
|  | ✓ | TG*_231630 | IMC4 | IMC (48) |
|  | ✓ | TG*_230210 | IMC10 | IMC (8, 74) |
|  | ✓ | TG*_235340 | ISC1 | IMC (64) |
|  | ✓ | TG*_252880 | CRMP | Conoid (75) |
|  | ✓ | TG*_246720† | Hypothetical | Conoid (48) |
| ✓ | ✓ | TG*_239885 | Prot. Kinase | Unknown (76) |
| **Localized in this study** |  |  |  |  |
| **Y2H** | **BioID** | **Gene ID** | **Gene Name** | **Localization** |
| ✓ | ✓ | TG*_238380 | Hypothetical | Apical cap |
| ✓ | ✓ | TG*_285140 | Hypothetical | Apical dot |
| ✓ |  | TG*_236060 | Hypothetical | Cytosolic/apical cap |
| ✓ |  | TG*_260500‡ | Hypothetical | IMC sutures |
|  | ✓ | TG*_231210‡ | Epsin-like | Apical/endosomal? |
|  | ✓ | TG*_309560‡ | Bax-like | Apical/endosomal? |
| ✓ |  | TG*_315950 | CSAR1 | Residual body (-IAA) / apical cap (+IAA) |

†Previously identified at conoid by proximity biotinylation (77)
‡Previously identified in published conoid proteome (48)

**Table 1: Top hits from the ERK7 interactome.**
| Known localization |  |  |  |  |
| --- | --- | --- | --- | --- |
| Y2H | BioID | Gene ID | Gene Name | Localization |
| ✓ | ✓ | TG*_246950 | AC9 | Apical cap (37, 38, 65) |
| ✓ | ✓ | TG*_292950 | AC10 | Apical cap (38, 46) |
|  | ✓ | TG*_250820 | AC2 | Apical cap (60) |
|  | ✓ | TG*_214880 | AC4 | Apical cap (60) |
|  | ✓ | TG*_229640 | AC8 | Apical cap (65) |
|  | ✓ | TG*_310030 | TgCAP | Apical cap (61, 66) |
|  | ✓ | TG*_310930 | TgFBXO1 | IMC (67) |
|  | ✓ | TG*_316540 | ISP3 | IMC (60) |
|  | ✓ | TG*_231640 | IMC1 | IMC (68, 69) |
|  | ✓ | TG*_231630 | IMC4 | IMC (45) |
|  | ✓ | TG*_230210 | IMC10 | IMC (8, 70) |
|  | ✓ | TG*_235340 | ISC1 | IMC (60) |
|  | ✓ | TG*_252880 | CRMP | Conoid (71) |
|  | ✓ | TG*_246720† | Hypothetical | Conoid (45) |
| ✓ | ✓ | TG*_239885 | Prot. Kinase | Unknown (72) |
| Localized in this study |  |  |  |  |
| Y2H | BioID | Gene ID | Gene Name | Localization |
| ✓ | ✓ | TG*_238380 | Hypothetical | Apical cap |
| ✓ | ✓ | TG*_285140 | Hypothetical | Apical dot |
| ✓ |  | TG*_236060 | Hypothetical | Cytosolic/apical cap |
| ✓ |  | TG*_260500‡ | Hypothetical | IMC sutures |
|  | ✓ | TG*_231210‡ | Epsin-like | Apical/endosomal? |
|  | ✓ | TG*_309560‡ | Bax-like | Apical/endosomal? |
| ✓ |  | TG*_315950 | CSAR1 | Residual body (-IAA) /<br>apical cap (+IAA) |

†Previously identified at conoid by proximity biotinylation (73)
‡Previously identified in published conoid proteome (45)

For yeast-two-hybrid, the full-length *Toxoplasma* ERK7 sequence (residues 2-692) was used as bait to probe a cDNA library derived from RH tachyzoites. From 267 individual clones, we identified 21 distinct candidate interactors (Table 1 and Supplemental Data S1; Figure S1), 13 of which we consider of highest quality based on the number of unique clones recovered for each. Notably, 8 of our yeast-two-hybrid hits were also identified by BioID, indicating good agreement between the two methods. Candidate ERK7 interactors were enriched in both known and newly identified proteins comprising IMC cytoskeleton, potentially involved in protein/membrane trafficking, and proteins that, like ERK7, concentrate at the apical end of the parasite during daughter budding (Table 1 and Supplemental Figure 1).

One candidate interactor identified in the Y2H screen stood out for the stark difference in its localization upon ERK7 degradation (Figure 2). We found that TG*_315950 had very low signal in mature parasites and at early stages of division (Figure 2A, Supplemental Figure S2). However, we observed high punctate signal restricted to maternal cytosol late in the budding process (Figure 2A, Supplemental Figures S2). Very late in budding, likely just before the parasites begin cytokinesis, TG*_315950 signal appears restricted to the residual body (Supplemental Figures S2). The TG*_315950 signal remains at the residual body throughout cytokinesis and in mature parasites that have completed division (Figure 2A, Supplemental Figure S2). Thus, TG*_315950 appears to have a role late in the budding process, though it is concentrated at the residual body. Unexpectedly, in parasites grown in IAA to induce ERK7 degradation, we found that TG*_315950 concentrates at the daughter apical caps and conoids late in the budding process (Figure 2B). Thus loss of ERK7 causes a mislocalization of TG*_315950 protein. Intriguingly, in normal parasites with functional ERK7, TG*_315950 does not concentrate at the apical cap with the kinase (Figure 2C), which we have previously shown to be the pool of ERK7 functionally relevant to conoid maturation (39, 49). This lack of colocalization, combined with the apparent turnover of the protein during early budding, likely explains why it was not enriched in our BioID dataset.

**Figure 2:**
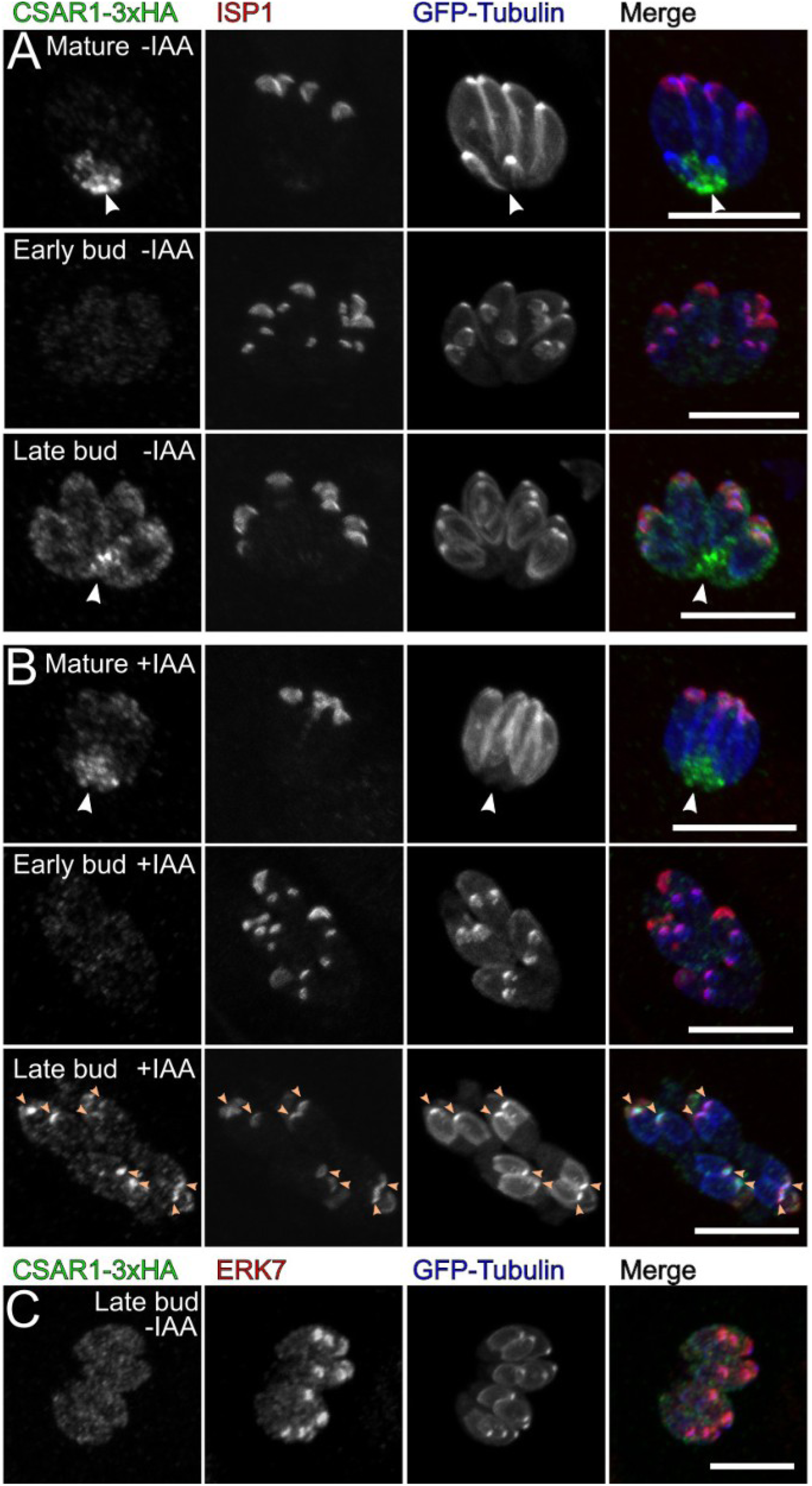
CSAR1 mislocalizes to daughter buds upon ERK7 degradation. Maximum intensity Z-projections of confocal stacks of CSAR1-3xHA ERK7AID parasites grown for 24 h (A) without IAA and (B) with IAA to degrade ERK7AID. Parasites were visualized with GFP-tubulin (blue), and antibodies against HA (green) and the apical cap marker ISP1 (red). Concentration of CSAR1 signal at the residual body is indicated with a white arrow. The apical ends of the late daughter buds in the +IAA condition are indicated with orange arrows. Parasites were captured at the indicated points in the cell cycle. (C) Maximum intensity Z-projects of confocal stack of CSAR1-3XHA ERK7AID parasites counterstained ith an anti-ERK7 antibody. All scale bars are 10 μm.

### Loss of CSAR1 allows parasites to complete the lytic cycle without ERK7, but is required for full parasite fitness

TG*_315950 encodes a 1822 residue protein with a predicted RING domain at its C-terminus, suggesting that it functions as an E3 Ub ligase. Given that TG*_315950 mislocalizes to the apical caps of budding daughters upon ERK7^AID^ degradation (Figure 2B), we reasoned that TG*_315950 may have aberrant function of its E3 ligase activity in which it inappropriately targets daughter conoids for degradation when ERK7 is not correctly localized to the daughter buds. For its apparent role in turnover of the parasite cytoskeleton (described below), we have named TG*_315950 CSAR1 (<u>C</u>ytoskeleton <u>Sa</u>lvaging <u>R</u>ING family member 1). If CSAR1 does indeed mediate the loss of the daughter conoids upon ERK7 loss-of-function, disruption of CSAR1 activity should rescue the ERK7 conoid phenotype. To address this question, we attempted to create a strain in which both ERK7 and CSAR1 were tagged with AID and therefore could be simultaneously inducibly degraded. We were unable to visualize any 3xHA signal of CSAR1 C-terminally tagged with AID-3xHA, and the parasites had a markedly reduced fitness, suggesting that we had produced either a hypomorph or null allele of CSAR1, in which the protein was constitutively degraded. We therefore used CRISPR/Cas9-assisted homologous recombination to replace the CSAR1 coding sequence with an HXGPRT selection cassette in the background of the ERK7^AID^(eGFP-tubulin) strain. This RH*Δcsar1* (in ERK7^AID^/eGFP-tubulin; hereafter, referred to as *Δcsar1*) strain fully reproduced the phenotype of the CSAR1^AID^ strain. Importantly, *Δcsar1* had no effect on auxin-mediated degradation of ERK7 protein (Figure 3A). We used a plaque assay to compare gross parasite fitness through the lytic cycle (Figure 3B,C). As expected, we observed no plaques from ERK7^AID^ parasites grown in the presence of IAA. In addition, we saw no loss of parasite fitness when wild-type CSAR1 was tagged with 3xHA, indicating the tag does not disrupt CSAR1 function (Figure 3B,C). Strikingly, *Δcsar1* parasites showed equivalent plaque efficiency whether or not ERK7 had been degraded (Figure 3B). In addition, the plaque efficiency was not significantly different from that of the parental ERK7^AID^ strain grown in the absence of IAA. These data demonstrate that the loss of CSAR1 suppresses the essentiality of ERK7 in the *Toxoplasma* lytic cycle.

**Figure 3:**
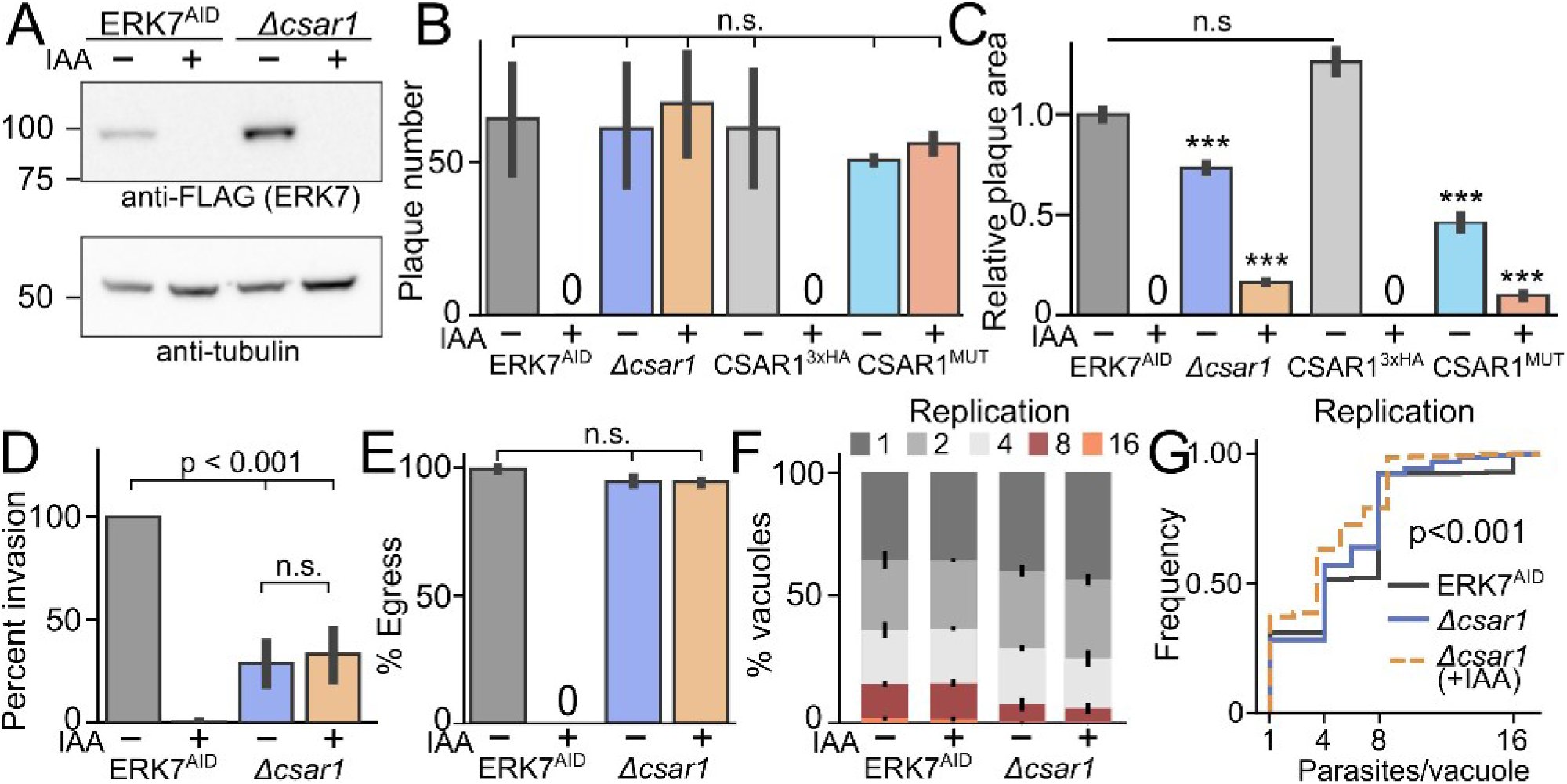
CSAR1 is required for full efficiency of lytic cycle. (A) Western blot stained with anti-FLAG (recognizing ERK7AID-3xFLAG) and anti-tubulin of lysates from ERK7AID and Δcsar1 parasites grown in ±IAA. Quantification of (B) plaque number, (C) relative plaque size, (D) invasion rate, (E) ionophore-induced egress, and (F) replication rates of ERK7AID or Δcsar1 parasites grown in ±IAA binned as indicated. (B,C) also show relative fitness of the wild-type 3xHA-tagged and RING-domain mutant CSAR1 (C1791A/H1792A/H1796A; CSAR1MUT) parasites. Significance on (B-E) calculated by 1-way ANOVA followed by Tukey’s multiple comparison test (*** is p<0.001; n.s., not significant). (G) is the unbinned cumulative frequency of the data from (F). Invasion, egress, and plaque assays are n=3 biological replicates conducted with n=3 technical replicates. Replication quantified from 3 biological replicates of n≥100 vacuoles. Error bars on (F) are s.d., all others are 95% confidence-interval of mean. Replication of Δcsar1 compared to parental ERK7AID is p<0.001 as calculated by Kolmogorov-Smirnov test.

While loss of CSAR1 did not reduce the plaquing efficiency as compared with the ERK7^AID^ parental strain, the plaque sizes were significantly smaller (Figure 3C). In the absence of IAA, *Δcsar1* parasites produced plaques ~70% the size of the parental ERK7^AID^ strain. In the presence of IAA, *Δcsar1* parasites produced plaques only ~15% the size of the parental strain in the −IAA condition, indicating that degradation of ERK7 further reduces fitness in these parasites, which may synergize with the loss of fitness caused by deletion of *Δcsar1*. The plaque number and size simultaneously informs on every step in the *Toxoplasma* lytic cycle: invasion, replication, and egress. We therefore tested the *Δcsar1* fitness for each of these steps.

To test invasion efficiency, we allowed 1×10^6^ parasites of either Δcsar1 or parental ERK7^AID^ strain (grown ±IAA for >12 hours prior to experiment) to invade for 1 hour before fixation. Consistent with our previously published observations (37, 39), degradation of ERK7 blocked attachment and invasion in the ERK7^AID/IAA^ parasites (Figure 3D). On the other hand, while *Δcsar1* parasites invaded less efficiently than the parental ERK7^AID^, their invasion was unaffected by growth in IAA (Figure 3D). Thus CSAR1 is required for the ERK7 phenotype on invasion.

We next tested whether deletion of CSAR1 altered the efficiency of parasite egress from an infected cell and the parasites’ motility, once extracellular. Parasites were allowed to invade fibroblasts for 4 h and then grown for 24-36 h in the presence or absence of IAA. Egress was induced by incubation with calcium ionophore for 1 min and parasites were immediately fixed. We quantified efficiency of egress by two metrics: by scoring whether the parasite vacuole had ruptured, using GRA1 as a marker for an intact vacuole (Figure 3E). As expected, all vacuoles of the ERK7^AID^ strain had egressed within 1 min when grown in the absence of IAA, but ERK7^AID/IAA^ parasites exhibited a complete block in egress (Figure 3E). The *Δcsar1* parasites, however, readily egressed even when grown in IAA (Figure 3D), demonstrating that loss of CSAR1 rescues the ERK7 egress phenotype, as well. However, we noted that *Δcsar1* parasites appeared to have a modest delay in egress, as ~5% of vacuoles remained intact during the treatment with ionophore, regardless of treatment with IAA. However, while this slight delay in egress was reproducible, it was not statistically significant (p > 0.05).

As a mild defect in egress was unlikely to be responsible for the reduction in plaque size we observed in the *Δcsar1* parasites, we reasoned that replication rate may be affected, as well. We therefore allowed parasites of both strains to invade for 4 h before changing the media to either ±IAA for 24 h growth. We stained parasites with anti-IMC1 antibody and Hoechst and quantified the number of parasites in each vacuole (Figure 3F,G). Consistent with their smaller plaques, we observed that *Δcsar1* parasites had a substantial reduction in their replication rate as compared with the ERK7 ^AID^ parental and this phenotype was further exacerbated by growth in IAA, indicating loss of ERK7 still reduces fitness in these parasites. Therefore, loss of CSAR1 suppresses the essentiality of ERK7 at the cost of efficient parasite replication.

### CSAR1 is required for recycling of maternal cytoskeleton during cytokinesis

To examine if there were any gross defects in the *Toxoplasma* cellular morphology in the *Δcsar1* parasites, we compared the parental ERK7^AID^ and *Δcsar1* parasites by fluorescence microscopy. Parasites were grown in fibroblasts for 18-24 h, fixed, and microtubule structures visualized with GFP-tubulin. We also stained with antibodies against ERK7 and IMC1, which we used as a marker for the inner membrane complex that outlines each parasite and growing buds (Figure 4A,B). Both ERK7^AID^ and *Δcsar1* parasites appear normal. However, *Δcsar1* vacuoles retain the apical complex and microtubule cytoskeleton in the parasite residual body after each replication. We observed n-1 retained maternal cytoskeletons per parasite in each *Δcsar1* vacuole (*e.g.* 1 in 2-pack, 3 in a 4-pack). Thus CSAR1 is essential for turnover and salvaging of these microtubule structures.

**Figure 4:**
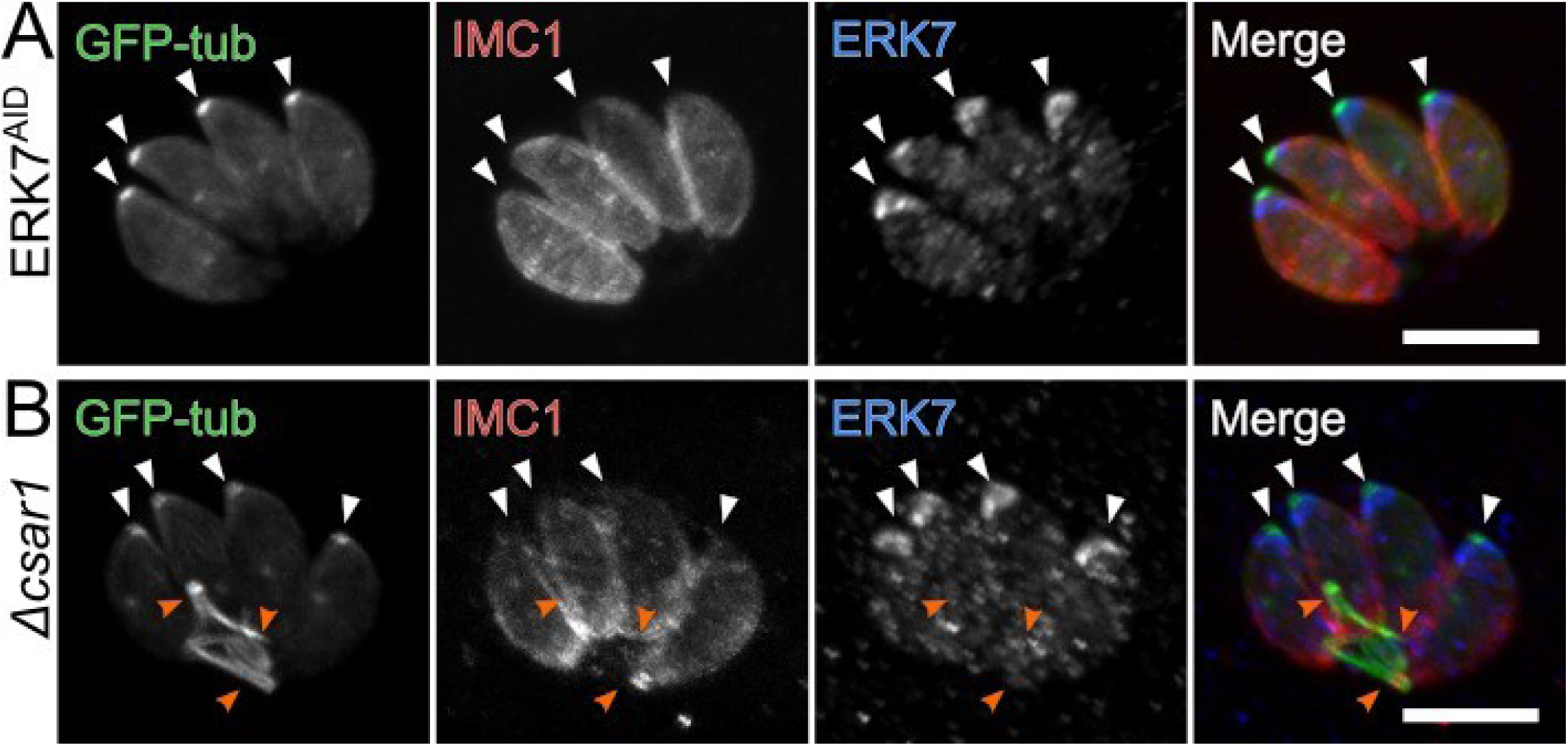
Loss of CSAR1 causes retention of maternal cytoskeleton in the residual body. Maximum intensity Z-projections of confocal stacks of (A) ERK7AID and (B) Δcsar1 parasites grown in the absence of IAA. Parasites were visualized with GFP-tubulin (green), and antibodies against the inner membrane complex marker IMC1 (red) and ERK7 (blue). Apical ends of parasites are indicated with white arrows. Retained cytoskeleton from the previous two divisions in the Δcsar1 vacuole is indicated with orange arrows. Scale bars are 5 μm.

We next sought to verify that the retained maternal cytoskeleton was, indeed, in the residual body. We examined *Δcsar1*-infected cells by correlated-fluorescence and transmission electron microscopy (Figure 5, Supplemental Figure S3). In vacuoles with *Δcsar* parasites, we observed residual body structures with clear connections to the parasites that contained extended filamentous projections consistent with retained microtubules, (Figure 5A-C) as well as retained conoids (Figure 5B-B’’). We also observed transverse slices through bundles of ~20 microtubules, consistent with retained maternal subpellicular microtubules (Figure 5D). In all cases, these cytoskeletal structures were clearly delineated by a single membrane, consistent with their retention in the residual body, which appears to be distorted to accommodate the length of the microtubule structures.

**Figure 5:**
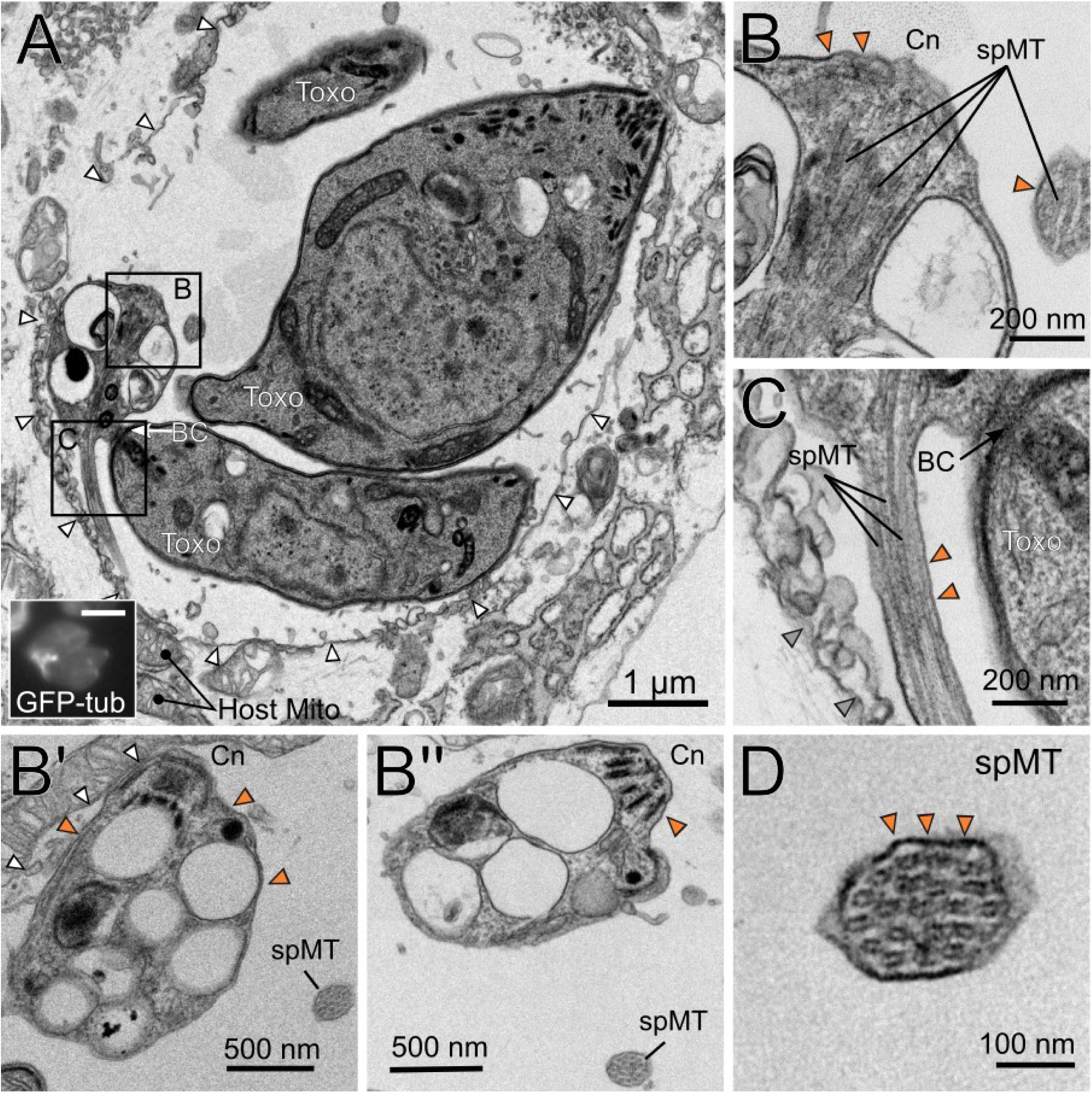
Retained maternal cytoskeleton is contained in membrane of the residual body. (A) Correlated fluorescence (GFP-tub; inset scale bar 5 μm) and transmission electron microscopy of *Δcsar1*-infected fibroblasts. Boxed regions are zoomed in the indicated panels. (B,C) Structures consistent with parasite microtubule cytoskeleton are found within the residual body. (B’,B’’) serial slices of the region in (B) show multiple retained conoids are tightly packed within the residual body. (D) Cross-section of a bundle of 20 subpellicular microtubules delineated by the residual body membrane. BC – basal complex; Cn – conoid; spMT – subpellicular microtubules; white arrowheads – parasitophorous vacuolar membrane; orange arrowheads – residual body membrane; Host Mito – host mitochondria. Images are representatitive of n=4 vacuoles imaged by TEM.

#### Loss of CSAR1 leads to increased plasticity in daughter cell number during Toxoplasma division

We reasoned that the blocked recycling of maternal microtubules in *Δcsar1* parasites may lead to a reduction in the availability of soluble tubulin. To test this, we compared the ratios of soluble to insoluble, polymerized, tubulin in *Δcsar1* and parental parasites that had been mechanically released from their host cells. This treatment also disrupts the residual bodies, allowing us to purify the intact parasites from the maternal cytoskeleton retained therein. We found that *Δcsar1* parasites had ~2-fold less soluble tubulin than the parental parasites (Figure 6A, Supplemental Figure S4A; 20±4% versus 9±3% of total). If there was, indeed, a reduced pool of soluble tubulin in these parasites, we reasoned that they would show increased sensitivity to drugs that target tubulin polymerization, such as oryzalin (50). Consistent with this idea, we found that *Δcsar1* parasites showed reduced viability (plaque number) at concentrations of oryzalin at which the paternal parasites were insensitive (Supplemental Figure S4B). Because microtubule dynamics are intimately coupled to the concentration of available soluble tubulin (51), and these dynamics are required for mitotic spindle formation, we reasoned it was possible that loss of CSAR1 may affect the efficiency of *Toxoplasma* replication, which would help explain its reduced growth rate (Figure 3F,G).

**Figure 6:**
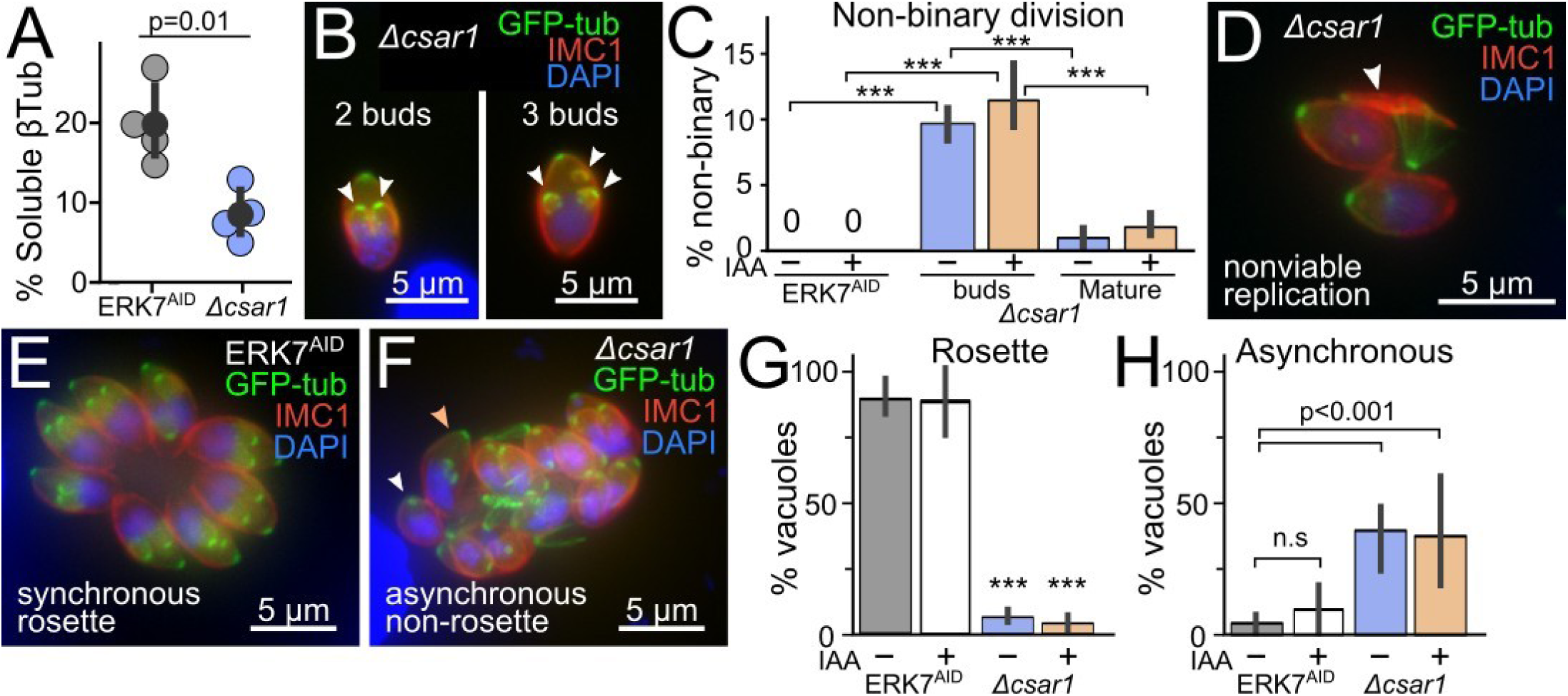
*Δcsar1* parasites exhibit multiple defects in cell cycle. (A) The percentage of soluble tubulin (versus membrane-insoluble assembled cytoskeleton) was quantified in n=4 biological replicates (averaged from n=4 technical replicates) of parental ERK7^AID^ versus *Δcsar1* parasites. Significance estimated by unpaired Student’s t-test. Western blots for these data are in Supplemental Figure S4. (B) Representative mages of *Δcsar1* parasites dividing normally (2 buds) or abnormally (3 buds). (C) Quantification of non-binary division for ERK7^AID^ and *Δcsar1* parasites. “buds” and “mature” respectively indicate actively dividing parasites or newly divided, mature parasites. (D) Representative image of a recently divided *Δcsar1* parasite showing the remains of a nonviable parasite (white arrow). Representative images of (E) ERK7^AID^ parasites dividing synchronously within a normal rosette vacuole and (F) *Δcsar1* parasites dividing asynchronously (white arrow: nonbudding; orange arrow: early buds; other parasites: late buds) in a non-rosette vacuole. Quantification of (G) Rosette/non-rosette and (H) Asynchronous division phenotypes. Phenotypes quantified from 3 biological replicates of n≥100 vacuoles. Error bars are 95% confidence interval of mean. p-values are 1-way ANOVA followed by Tukey’s test. (***, p < 0.001)

*Toxoplasma* replicates by endodyogeny, in which two daughter buds mature within the maternal cell (Figure 1). Normal division involves the generation of exactly two daughter buds. however, in rare cases (<1%), wild-type *Toxoplasma* division can result in 3 or more buds (17, 52). In imaging cells to examine replication, we noticed that *Δcsar1* parasites appeared to have an unusually high number of aberrant budding events (Figure 3G, Figure 6B). To compare the extent of non-binary parasite budding in *Δcsar1* versus the parental strain, we allowed parasites to invade for 2 h, washed off extracellular parasites, and allowed parasites to grow for 6 h at 37°C before being fixed and stained for imaging. To test the effect of ERK7 degradation on aberrant parasite budding, parasites were grown in the presence or absence of IAA. We quantified the number of buds per parasite for each condition (Figure 6C). While we did not observe aberrant dividing ERK7^AID^ parasites, a remarkable 10±1% of *Δcsar1* parasites had >2 buds per division (Figure 6C). This phenotype was not further exacerbated by growth in IAA, indicating loss of ERK7 does not affect the daughter-count during budding.

A previous analysis of rare non-binary division in wild-type *Toxoplasma* indicated that the majority of parasites resulting from non-binary division were viable (52). Because the maternal cytoskeleton is retained in the residual body of *Δcsar1* parasites (Figures 4,5), we can unambiguously identify parasites that have divided exactly once since infection. We used this fact to test whether non-binary division events produced viable parasites in *Δcsar1*. We defined viable parasites as objects that showed appropriate signal for each GFP-tubulin, IMC1, and DAPI. After a single round of division, we found that only 1±0.7% vacuoles contained more than 2 mature parasites, which is substantially lower than expected by the ~10% of non-binary buds we observe in actively dividing parasites. Instead, we observed that many vacuoles contained what appeared to be malformed parasites in their residual bodies, lacking either a complete nucleus, inner membrane complex, or microtubule cytoskeleton (Figure 6D). Thus it appears that, unlike what has been reported for wild-type parasites (52), many of the non-binary division events in *Δcsar1* produce non-viable progeny.

Normally*, Toxoplasma* divide to form a rosette within a vacuole, such that the apical ends are oriented away from the residual body at the vacuole’s center. Furthermore, normal parasite division is synchronized within a vacuole. This synchrony has been attributed to active trafficking of cytosolic components between parasites along a network of actin filaments that is organized within the residual body (13, 14). Given that CSAR1 localizes to the residual body (Figure 2) and its disruption causes aberrant division (Figure 6B-D), we reasoned that disruption of CSAR1 may also affect vacuole organization and synchrony of division. To this end, we analyzed the images we had used to quantify parasite replication for these phenotypes. 90±5% of ERK7^AID^ vacuoles showed typical rosette arrangement regardless of IAA treatment and 90±10% these vacuoles were dividing synchronously (Figure 6E-H). In contrast, 94±1% of *Δcsar1* vacuoles had lost the rosette organization, though degradation of ERK7 with addition of IAA did not further impact the phenotype (Figure 6G-H). Similarly, while all parental vacuoles were dividing synchronously, we observed that 40±10% *Δcsar1* vacuoles contained parasites at markedly different points in the cell cycle (Figure 6G-H).

#### In absence of ERK7, CSAR1 mediates degradation of maturing daughter conoids

Disruption of CSAR1 enabled ERK7^AID^ parasites to escape the block in the lytic cycle due to ERK7 degradation (Figure 3B). Loss of CSAR1 also blocks turnover of maternal apical complex and microtubule cytoskeleton after division (Figure 4) and CSAR1 mislocalizes to the daughter apical caps upon ERK7 degradation (Figure 2B). Taken together with its predicted function as an E3 ubiquitin ligase, these data suggested that CSAR1 was responsible for the loss of daughter conoids in parasites in which ERK7 had been degraded. We used confocal microscopy to assess this model. Consistent with published data (37), we observed normal conoids in ERK7^AID^ parasites, though the conoid was lost in these parasites upon ERK7 degradation (Figure 7B). In the *Δcsar1* parasites, however, not only is does the maternal cytoskeleton build up in the residual body after replication (Figure 7C), but we also observe strong tubulin puncta in all parasites after treatment with IAA, consistent with preservation of the conoid (Figure 7D).

**Figure 7:**
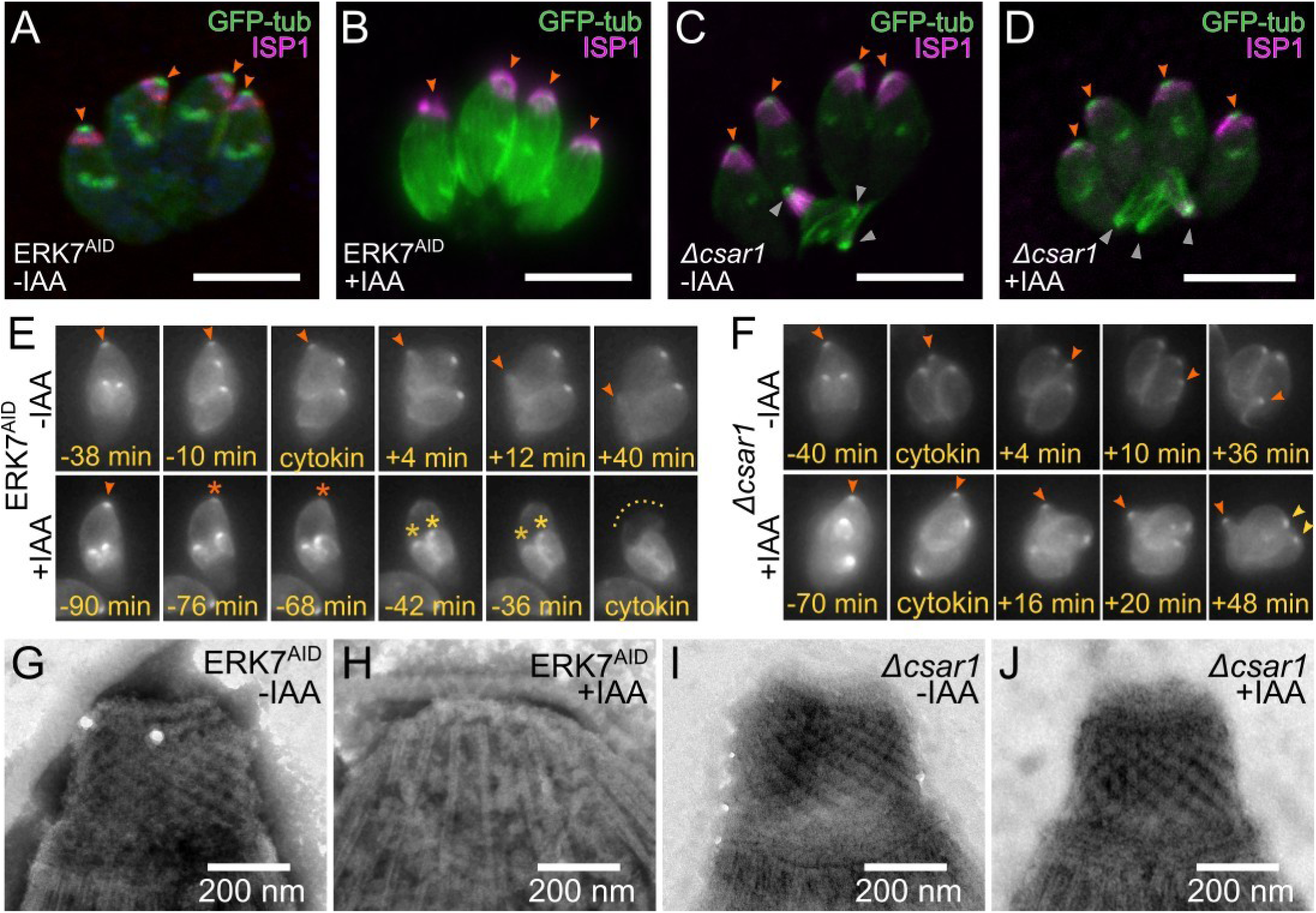
Loss of CSAR1 protects the conoid upon ERK7 degradation. (A-D) Maximum intensity Z-projections of confocal stacks of the indicated parasite strains grown in ±IAA and visualized with GFP-tubulin (green) and an antibody against the apical cap marker ISP1 (magenta). Apical ends of parasites are indicated with orange arrows. Retained cytoskeleton in the *Δcsar1* residual body is indicated with gray arrows. All scale bars are 5 μm. (E) Stills from live cell imaging of GFP-tubulin in dividing ERK7^AID^ parasites grown in ±IAA. The time before (-) and after (+) cytokinesis is indicated. Maternal conoid is indicated with an orange arrow (or orange star when degraded). Daughter conoids are indicated with gold stars at point of degradation. (F) Stills from *Δcsar1* parasites imaged as in (E). Annotations as above; yellow arrows indicate viable daughter conoids at end of division in +IAA media. (G-I) Negative-stained TEM of detergent-extracted cytoskeleton “ghosts” from the indicated strains and grown in ±IAA. (Representative of n≥30 TEM images).

To better understand the timing of conoid loss upon ERK7 degradation, we used video microscopy of eGFP-tubulin signal to follow parasite budding and cytokinesis in living ERK7AID and Δcsar1 parasites grown in IAA. In ERK7AID parasites grown in IAA, we observed that the maternal conoid is lost 78±47 min before the initiation of cytokinesis (Figure 7E, Supplemental Movie 1), and the daughter conoid signal is lost 31±12 min later (n=17 parasites). The loss of the maternal conoid in the first cycle of division after ERK7 degradation is inconsistent with our original model that ERK7 is required for assembly of a functional conoid (37), and instead suggests that the mature conoid is degraded in these parasites. Consistent the preservation of the conoid we observe by confocal imaging (compare Figures 7A,B to 7C,D), we see no effect on the GFP-tubulin conoid signal during division of *Δcsar1* parasites grown in IAA (Figure 7F, Supplemental Movie 1).

To test whether there are obvious changes in the apical complex ultrastructure in *Δcsar1* parasites as compared to the parental ERK7^AID^ strain, we visualized detergent-extracted parasite cytoskeleton ghosts by transmission electron microscopy (Figure 7G-J). As expected, the parasite cytoskeleton appears normal in ERK7^AID^ parasites untreated with IAA (Figure 7G), but the conoid is completely missing after growth in IAA (Figure 7H). In *Δcsar1* parasites, all detergent-stable cytoskeleton structures appeared normal, regardless of whether ERK7 had been degraded with IAA (Figure 7I-J). Thus deletion of *Δcsar1* does not appear to negatively impact apical complex assembly or function, and appears to protect the overall structure of the conoid when ERK7 has been degraded (Figures 7J).

CSAR1 is a putative RING-family E3 ubiquitin ligase. To test whether the CSAR1 predicted RING domain was required for its function, we created parasites in which the CSAR1 RING domain had been inactivated by mutating predicted Zn-binding sites (C1791A/H1792A/H1796A; Supplemental Figure S5) (53). The mutant CSAR1 (CSAR1^MUT^) was expressed in frame with a C-terminal 3xHA tag in the background of the ERK7^AID^ strain and is otherwise isogenic with the CSAR1^3xHA^ strain. Consistent with a role as an active E3 ligase, the CSAR1^MUT^ parasites phenocopied *Δcsar1*, as we observed retained maternal cytoskeleton in the residual bodies of replicating parasites (Figure 8). CSAR1^MUT^ showed similar cell cycle dependence to its localization as did the wild-type CSAR1^3xHA^ (Figure 2A), and concentrated in the residual body of mature parasites when ERK7 was present (Figure 8A). Strikingly, upon degradation of ERK7 by growth in IAA, CSAR1^MUT^ relocalized to both maternal and daughter apical caps of late budding parasites, though parasites did not lose their conoids (Figure 8B). Thus mutation of the CSAR1 RING domain, like complete knockout of the protein, suppresses ERK7 degradation, and allows parasites to complete the lytic cycle (Figure 3B).

**Figure 8:**
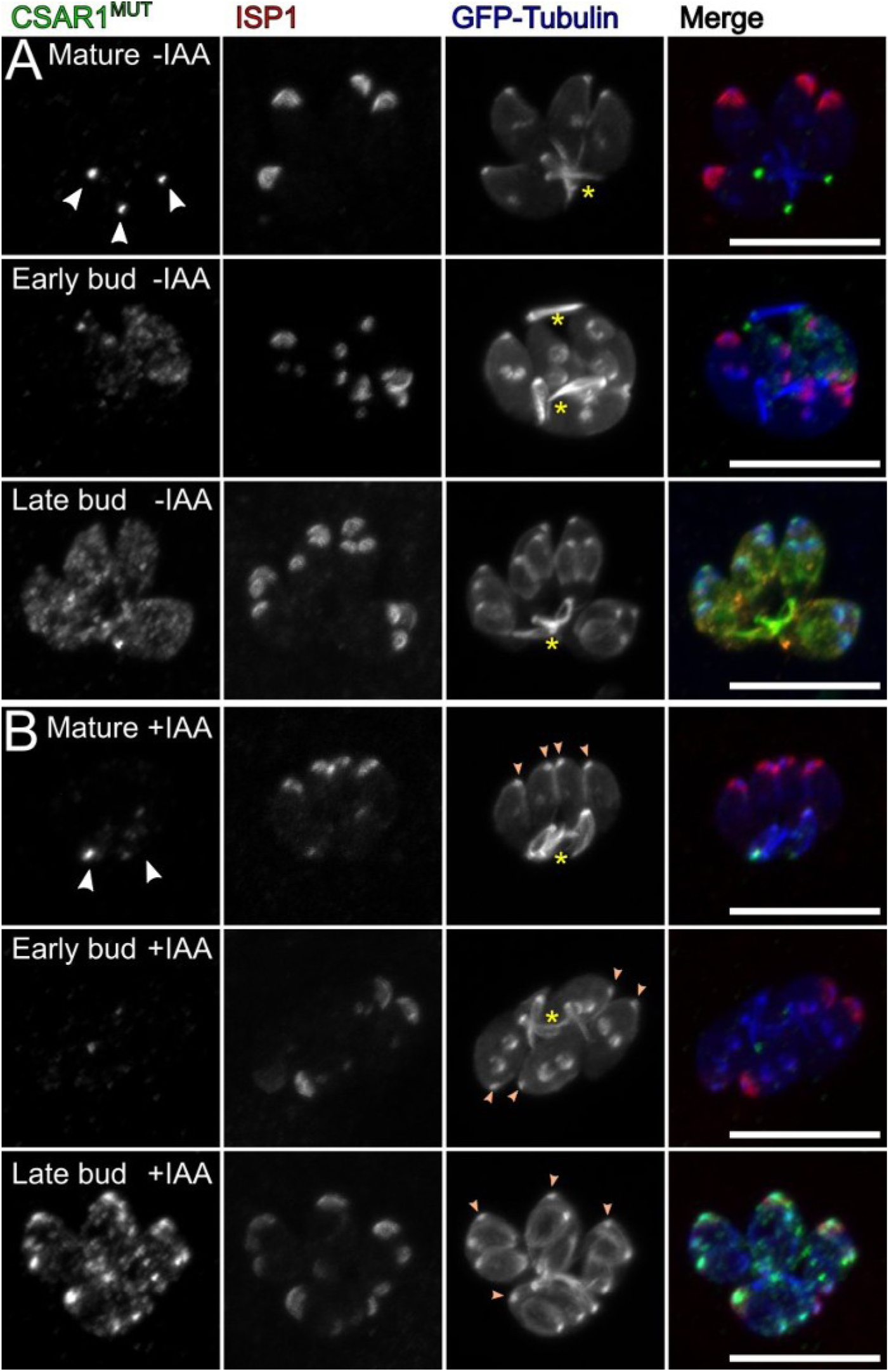
CSAR1 RING domain is required for function. Maximum intensity Z-projections of confocal stacks of CSAR1^MUT^-3xHA (ERK7^AID^) parasites grown for 24 h (A) without IAA and (B) with IAA to degrade ERK7^AID^. Parasites were visualized with GFP-tubulin (blue), and antibodies against HA (green) and the apical cap marker ISP1 (red). Note that CSAR1^MUT^ signal concentrates with retained cytoskeleton (yellow stars) in the residual bodies of mature parasites (white arrowheads) regardless of whether ERK7 is present. Also note that CSAR1^MUT^ parasites retain their maternal conoids (orange arrowheads) after replication, even when ERK7 has been degraded.

Taken together, these data support the model that CSAR1 is an E3 ubiquitin ligase that is essential for the turnover of the maternal cytoskeleton during endodyogeny, and that its subcellular localization, and therefore substrate targeting, is tightly regulated by ERK7 (Figure 9).

**Figure 9:**
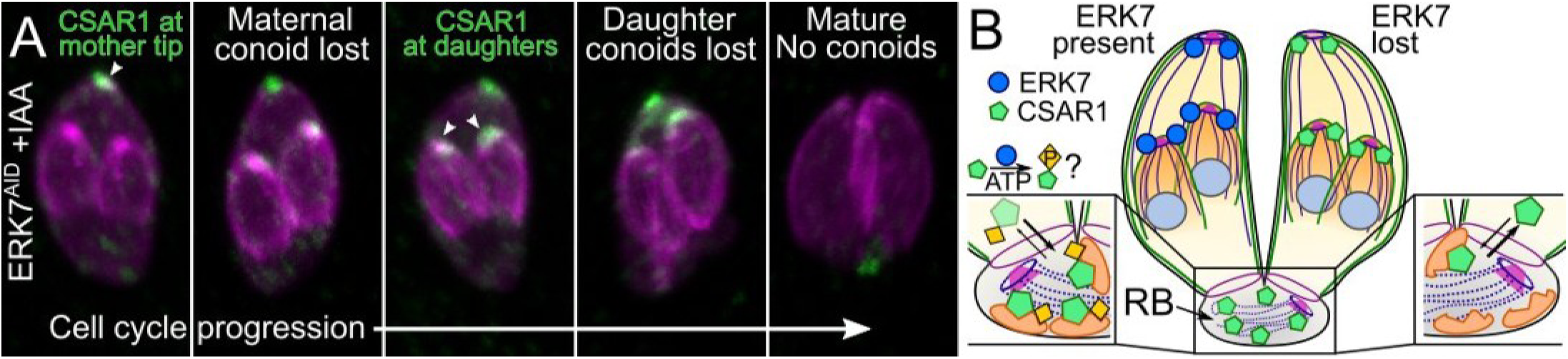
Model of functional relationship between CSAR1 and ERK7. (A) Pseudo-timecourse of cell cycle using fixed confocal micrographs of CSAR13xHA(ERK7AID) parasites grown in +IAA for 5 h. CSAR1 signal appears to build up first at the maternal and then at the daughter conoids just before the loss of the structures. (B) In normal parasites, ERK7 concentrates at the apical caps of both maternal and daughter parasites and drives CSAR1 to concentrate in the residual body (RB), possibly through ERK7 kinase activity. This protects the maternal and daughter apical complexes from degradation. When ERK7 is lost, CSAR1 is not retained in the residual body and concentrates at the apical tips of mother and daughter cells, leading to premature degradation of the conoids.

## Discussion

We have delineated the protein interactome for *Toxoplasma* ERK7. In other organisms, ERK7 orthologs in have been associated with the regulation of variety of cellular processes, including ciliogenesis (54–56), protein trafficking and signaling (57–60), and autophagy (61). Because ERK7 appears to be found exclusively yet consistently in eukaryotes with ciliated cells (37, 41), it seems likely that its role in cilia biogenesis and maintenance is a primary conserved function. Consistent with this idea, our ERK7 interactome is enriched in proteins localizing to the parasite cytoskeleton and apical complex (Table 1). Intriguingly, among the putative interactors we identified for *Toxoplasma* include the autophagy processing enzyme ATG4 and a number of proteins that appear to localize to the parasite endolysosomal trafficking system (Table 1, Supplemental Table S1, Supplemental Figure S1). These findings suggest that other reported functions for ERK7 may be present in *Toxoplasma*, as well. Defining the relationship between ERK7 and these pathways in *Toxoplasma* and other organisms will determine whether there is, indeed, a core conserved ERK7 signaling pathway. We originally identified ERK7 as essential to the maturation of the *Toxoplasma* apical complex (37), and initiated this study to determine the downstream effectors that potentiate this function. We identified a putative RING-family E3 ligase, CSAR1, as this molecule. While we had originally envisioned the ERK7 loss-of-function phenotype as a loss of a positive regulation, we found that CSAR1 appears to function aberrantly upon ERK7^AID^ degradation. We found that CSAR1 is required for the turnover of the hyperstable maternal microtubule cytoskeleton and apical complex late in parasite division (Figure 4). Upon ERK7 degradation, CSAR1 mislocalizes to the late daughter bud apical cap and conoid (Figures 2, 10) shortly before their disappearance (Figure 7, Supplemental Movie 1). Consistent with a function as an E3 ligase targeting the parasite cytoskeleton, *Δcsar1* parasites in the ERK7^AID^ background completely rescue the loss of conoid phenotype upon ERK7 degradation. Importantly, we identified CSAR1 as an ERK7 interactor by yeast-two-hybrid (Table 1), indicating that these proteins directly interact. Taken together, these data lead us to a model where ERK7 localizes to the apical caps of developing daughter parasites both to promote trafficking and assembly of components of the apical complex and IMC cytoskeleton and to protect the daughter buds from premature degradation by CSAR1-mediated ubiquitination (Figure 9).

It has long been appreciated that turnover of maternal secretory organelles (4) and cytoskeleton (7, 17, 62) was a visible step in parasite division. Furthermore, this process has been suggested to occur at the residual body since the organelle’s identification by electron microscopy over 60 years ago (10, 11). However, to date, very few proteins that natively and functionally localize to residual body have been identified. We have demonstrated that CSAR1 is involved in protein turnover and normally concentrates at the residual body. This finding provides an important functional validation of the long-held hypothesis that the residual body is a site of organellar recycling. We found that CSAR1 levels begin to build up in the residual body and maternal cytosol late in daughter budding and become restricted to the residual body after cytokinesis. In addition to a retention of maternal cytoskeleton in the residual bodies of *Δcsar1* vacuoles (Figure 4), we observed disruption of parasite organization within the vacuole and dysregulation of synchronicity of cell cycle and of daughter-cell counting during replication (Figure 6). Similar phenotypes have been observed both with disruption of components of parasite IMC cytoskeleton (63, 64) and of the actin organization of the residual body (13, 14, 65). Notably, one of these proteins, TgCAP (65), is apically localized and was identified in our ERK7 interactome by BioID (Table 1). Whether the collection of *Δcsar1* phenotypes is a result of a physical block of the residual body by retained maternal cytoskeleton, dysregulation of *e.g.* monomeric tubulin levels, or disruption of an unidentified CSAR1 client remains to be determined.

An important facet of endodyogeny is that the maternal parasite is infectious until very late in the cell cycle; Toxoplasma may have evolved to divide by this process to maximize its infectivity. However, once the maternal secretory organelles begin to break down, which occurs late in budding (4), the parasite would lose its ability to invade. Thus the maternal cytosol may, at this point, already be considered to be functionally residual body and CSAR1 is an early marker of this transition. The normal versus aberrant functions of CSAR1 suggests Toxoplasma uses residual body localization to compartmentalize processes with potentially catastrophic consequences to cell health, and ERK7 is one regulator of this spatial restriction (Figure 9). Such a model argues the residual body should be considered a distinct and important organelle with a much more active role in the Toxoplasma lytic cycle than previously appreciated.

## Materials and Methods

*PCR and plasmid generation* – All PCR for plasmid generation was conducted using Phusion polymerase (NEB) using primers listed in Supplemental Table 1. Constructs were assembled using Gibson master mix (NEB).

*Parasite culture and transfection* – Human foreskin fibroblasts (HFF) were grown in Dulbecco’s modified Eagle’s medium supplemented with 10% fetal bovine serum and 2 mM glutamine. *Toxoplasma* tachyzoites were maintained in confluent monolayers of HFF. ERK7-BioID2-3xHA was created by transfecting the RH*Δku80Δhxgprt* strain (66) with 50 μL of a PCR product using Q5 polymerase (NEB) with 500 bp homology arms flanking a BioID2-3xHA tag together with 5 μg of a Cas9 plasmid that had been modified to express HXGPRT and also a gRNA targeting the C-terminus of ERK7 (37). The parasites were selected for HXGPRT expression for 2 days and immediately single cell cloned without selection. (ERK7p)mVenus-BioID2-3xHA was created by transfecting RH*Δhxgprt* with a plasmid containing mVenus in frame with BioID2-3xHA and driven by the ERK7 promoter (1011 bp upstream of the ERK7 start). 3xHA tagged interactome candidates were tagged in the background of the ERK7^AID^ strain (37) by transfecting 15 μg of linearized plasmids containing a ~1000 bp of targeting sequence in-frame with a C-terminal 3xHA tag. *Δcsar1* parasites were created by transfecting 5 μg of a plasmid expressing a Cas9 and a gRNA targeting the ATG of CSAR1 and 50 μL of a Q5 PCR product containing a floxed HXGPRT selection cassette flanked by homology arms 5’ and 3’ of the CSAR1 gene. All transfectants were selected with CDMEM + MPA/Xanthine and single cell cloned by serial dilution. All experiments were conducted with parasites that had been cultured for <1 month after cloning.

*Proximity biotinylation and mass spectrometry –* 10x 15 cm dishes of HFF were infected with either ERK7-BioID2-3xHA or with (ERK7p)mVenus-BioID2-3xHA and allowed to grow in the presence of 150 μM biotin until parasites began lysing the monolayer (~36 h). Parasites were released from host cells by passage through a 25G needle and washed with PBS twice. Parasites were incubated in 2.5 mL for 1 h at 4 °C with RIPA buffer supplemented with protease inhibitors (Sigma). To remove exogenous biotin, each sample was buffer exchanged a PD-10 desalting column (GE Healthcare) that had been equilibrated with RIPA buffer. Biotinylated proteins were separated on magnetic streptavidin resin (NEB) and eluted with 2× SDS loading buffer supplemented with 4% SDS and 10 mM Β-mercaptoethanol. Proteins were separated briefly on a pre-cast 4-20% SDS PAGE (BioRad) for 6.5 min at 200 V. The region of the gel containing the samples (~1 × 0.5 cm) was excised for downstream processing for mass spectrometry.

Samples were digested overnight with trypsin (Pierce) following reduction and alkylation with DTT and iodoacetamide (Sigma-Aldrich). The samples then underwent solid-phase extraction cleanup with an Oasis HLB plate (Waters) and the resulting samples were injected onto an Orbitrap Fusion Lumos mass spectrometer coupled to an Ultimate 3000 RSLC-Nano liquid chromatography system. Samples were injected onto a 75 μm i.d., 75-cm long EasySpray column (Thermo) and eluted with a gradient from 0-28% buffer B over 90 min. Buffer A contained 2% (v/v) ACN and 0.1% formic acid in water, and buffer B contained 80% (v/v) ACN, 10% (v/v) trifluoroethanol, and 0.1% formic acid in water. The mass spectrometer operated in positive ion mode with a source voltage of 1.8-2.4 kV and an ion transfer tube temperature of 275 °C. MS scans were acquired at 120,000 resolution in the Orbitrap and up to 10 MS/MS spectra were obtained in the ion trap for each full spectrum acquired using higher-energy collisional dissociation (HCD) for ions with charges 2-7. Dynamic exclusion was set for 25 s after an ion was selected for fragmentation.

Raw MS data files were analyzed using Proteome Discoverer v2.4 (Thermo), with peptide identification performed using Sequest HT searching against the *Toxoplasma gondii* and human protein databases from UniProt. Fragment and precursor tolerances of 10 ppm and 0.6 Da were specified, and three missed cleavages were allowed. Carbamidomethylation of Cys was set as a fixed modification, with oxidation of Met set as a variable modification. The false-discovery rate (FDR) cutoff was 1% for all peptides. Data have been deposited in the MassIVE database (MSV000088501).

*Yeast-2-hybrid screening* – Yeast two-hybrid screening was performed by Hybrigenics Services, S.A.S., Evry, France. The coding sequence for Toxoplasma gondii ERK7 was PCR-amplified and cloned into pB27 as a C-terminal fusion to LexA (LexA-ERK7). The construct was used as a bait to screen a random-primed Toxoplasma gondii wt RH strain library constructed into pP6. pB27 and pP6 derive from the original pBTM116 (Vojtek and Hollenberg, 1995) and pGADGH (Bartel et al., 1993) plasmids, respectively. 75 million clones (8-fold the complexity of the library) were screened using a mating approach with YHGX13 (Y187 ade2-101::loxP-kanMX-loxP, matα) and L40ΔGal4 (mata) yeast strains as previously described (Fromont-Racine et al., 1997). 267 His+ colonies were selected on a medium lacking tryptophan, leucine and histidine, and supplemented with 5 mM 3-aminotriazole, a competitive inhibitor of the HIS3 gene product, as we found that ERK7 exhibited mild autoactivation.. The prey fragments of the positive clones were amplified by PCR and sequenced to identify the corresponding interacting proteins.

*Plaque assays* – To measure plaque efficiency, 200 of each ERK7^AID^ and *Δcsar1* parasites were allowed to infect confluent HFF in one well of a 6 well plate in the either the presence or absence of IAA. After 7 days, the monolayer was fixed with MeOH, stained with crystal violet, and the resulting plaques counted. All plaque assays were performed in technical and biological triplicate. Plaque areas were quantified in ImageJ (67).

*Immunofluorescence* – HFF cells were grown on coverslips in 24-well plates until confluent and were infected with parasites. The cells were rinsed twice with phosphate buffered saline (PBS), and were fixed with 4% paraformaldehyde/4% sucrose in PBS at room temperature for 15 min. After two washes with PBS, cells were permeabilized with 0.1% Triton-X-100 in PBS for 10 min and washed 3x with PBS. After blocking in PBS + 3% BSA for 30 min, cells were incubated in primary antibody in blocking solution overnight at room temperature. Cells were then washed 3x with PBS and incubated with Alexa-fluor conjugated secondary antibodies (Molecular Probes) for 2 h and Hoechst, where appropriate. Cells were then washed 3x with PBS and then mounted with mounting medium (Vector Laboratories). Cells were imaged on either a Nikon A1 laser scanning confocal microscope with a 60x oil immersion 1.42 NA objective or with a Nikon Ti2E wide-field microscope with a 100x oil immersion 1.45 NA objective. Primary antibodies used in this study include rat anti-HA (Sigma; 1:1000 dilution), mouse m2 anti-FLAG (1:1,000 dilution; Sigma), rabbit anti-Tg-β-tubulin (1:10,000 dilution), guinea pig anti-TgERK7 (1:10,000 dilution), mouse anti-IMC1 (1:1000 dilution; gift of Gary Ward), anti-GRA1 (1:1000 dilution; BioVision), mouse anti-ROP2 (1:1000 dilution), mouse anti-MIC2 (1:1000 dilution; gift of Vern Carruthers), and mouse anti-ISP1 (1:1000 dilution; gift of Peter Bradley). Unless otherwise noted, micrographs are representative of 20 of 20 images collected.

*Live cell imaging* – ERK7^AID^ or *Δcsar1* parasites were syringe released from a T25 of a highly infected monolayer and passed through a 5 μm syringe filter (Pall) to remove cell debris. ~1×10^6^ parasites were added to a confluent monolayer of HFFs grown on a Lab-Tek 8-well #1.5 glass-bottomed chamber slide (Thermo) and allowed to invade and grow for 4-5 h at 37 °C in a 5% CO 2 incubator, after which time the well was washed extensively with warm HBSS (>5×) to remove extracellular parasites and the media was changed to either CDMEM ±500 μM IAA. The chamber slide was transferred to a Tokai Hit environment chamber mounted on a Ti2 Nikon wide-field microscope and maintained at 37 °C and 5% CO2. Parasite division was visualized by eGFP-α-tubulin fluorescence using a 100× NA 1.45 Pan Apo oil immersion objective (Nikon). Briefly, 10-15 regions were selected containing parasites with visible early buds. Images were collected every 2 min for 2-3 h as 10 μm stacks of 1 μm slices that were then maximum-intensity Z-projected. Movies were rendered using a custom Python script interfacing with Inkscape v1.1.1 and ffmpeg.

Western blotting – Proteins were separated by SDS-PAGE and transferred to a PVDF membrane. Membranes were blocked for 1 h in PBS + 5% milk, followed by overnight incubation at 4°C with primary antibody in blocking solution. The next day, membranes were washed 3× with TBST, followed by incubation at room temperature for 1-2 h with HRP-conjugated secondary antibody (Sigma) in blocking buffer. After 3× washes with TBST, western blots were imaged using ECL Plus reagent (Pierce) on a GE ImageQuant LAS4000. Antibodies used in this study include: Rb anti-Tg-β-tubulin (1:5,000 dilution) and mouse m2 anti-FLAG (Sigma; 1:1,000 dilution).

*Tubulin solubility assay* – T25 flasks of highly infected or freshly lysing Δcsar1 and ERK7^AID^ parasites were scraped and then mechanically released by passing through a 27 g syringe 5 times. The parasites were then passed through a 5 μm PVDF syringe filter. The cells were then pelleted at 250 g for 10 min. 5 ml of cold PBS was added to wash. 10 μl of parasites were placed on a hemocytometer to be counted. The parasites were pelleted again at 250 g for 10 min. The cell pellet was then brought up in 100 μl of a cold lysis buffer (1% Triton-100, 50 mM Tris pH 7.5, 150 mM NaCl). Working concentrations of E64, TAME, Pepstatin, and PMSF were added to buffer immediately prior to lysis. The parasites were agitated for 10min at 4° C and then pelleted at 16.3 k-g at 4°C for 30 min to separate the insoluble cytoskeleton from soluble protein. The entire soluble fraction was moved to a new microfuge tube and the insoluble pellet was brought up in cold 6 M urea. 20 μl of 6x SDS loading dye with BME were added to each tube. The samples were then sonicated in bath sonicator for 10 minutes, and then heated at 95°C for 10 minutes. SDS-Page gels were loaded with volumes normalized by parasite number in sample. Equal volume of pellet and supernatant samples were used. Four biological replicates for each sample with four technical replicates were used for each sample. The protein was transferred for western blot at 100V for 6hrs at 4°C. Membranes were blotted with Rb anti-Tg-β-tubulin. The western blots were quantified using the gel analyses function in ImageJ. *Invasion and egress assays* – For invasion assays, highly infected monolayers were mechanically disrupted by passage through a 27 gauge needle to release them. 1×10^6^ parasites of each strain tested (all expressing eGFP-α-tubulin) were added to HFF monolayer grown in a 24 well with coverslip and incubated for 2 h at 37°C. These were then washed 10× with PBS and fixed and prepared for imaging. 3.15 mm^2^ area (~1% of total well) were imaged per experiment and images analyzed in ImageJ. Invasion rates of all strains were normalized to the average of the ERK7^AID^ −IAA condition. For egress assays, parasites were allowed to grow for 24-36 h in HFF grown in a 24 well with coverslip. Parasites were incubated in pre-warmed HBSS containing 1 μM calcium ionophore A23187 (Cayman Chemical) for 1 min at 37°C before fixation in PFA, staining with anti-GRA1 antibody as a marker for intact vacuoles, and subsequent imaging. Egress rates were quantified by normalizing the number of intact vacuoles to a matching experimental set that was fixed without ionophore treatment.

*Transmission electron microscopy* – For negative stain EM of detergent extracted cytoskeletons: Parasite ghosts were prepared essentially as described in (68). T25 flasks of parasites were grown in either 500 μl IAA or vehicle media for 48 h. The highly infected T25s were syringe released, passed through a 5 μm filter, washed with Hanks Buffered Saline solution, and then brought up in 50 ul of 20 μM calcium ionophore in HBSS. The parasites were then incubated at 37 °C for 10 min. 4 μl of the parasite suspension were allowed to adhere to a grid, after which membranes were extracted by addition of 0.5% Triton-X-100 in PBS for 3-4 min. The grids were then briefly washed with MilliQ and then stained with 0.5% phosphotungstic acid, pH 7.4 for 30 s. The phosphotungstic acid was wicked off and the grids were again washed with MilliQ and allowed to dry. All TEM images were acquired on a Tecnai G2 spirit transmission electron microscope (FEI) equipped with a LaB6 source at 120 kV. For Correlated-fluorescence/electron microscopy: HFFs were plated on a MatTek 35mm dish with gridded coverslip. After 24 h the dishes were infected with Δ*csar1* parasites at an MOI of 3 into the low confluency HFF monolayer. Parasites were incubated at 37° C for 30hrs. Parasites were then imaged by DIC and fluorescence with a Nikon Ti2E wide-field microscope using an air 40x 0.95 NA objective to identify regions for sectioning just prior to fixation. The dish was then washed with PBS and fixed with 2.5% glutaraldehyde in 0.1M Na Cacodylate buffer plus 3% sucrose. After three rinses in 0.1 M sodium cacodylate buffer, the cells were postfixed with 1% osmium tetroxide and 0.8% K3[Fe(CN)6] in 0.1 M sodium cacodylate buffer for 1 h at room temperature. Cells were rinsed with water and en bloc stained with 2% aqueous uranyl acetate overnight. After three rinses with water, specimens were dehydrated with increasing concentrations of ethanol, infiltrated with Embed-812 resin, and polymerized in a 70°C oven overnight. Blocks were sectioned with a diamond knife (Diatome) on a Leica Ultracut UC7 ultramicrotome (Leica Microsystems) and collected onto copper grids poststained with 2% uranyl acetate in water and lead citrate. All TEM images were acquired on a Tecnai G2 Spirit transmission electron microscope (FEI) equipped with a LaB6 source at 120 kV. Alternating sections (between those mouted on grids) were mounted on slides and were then imaged by DIC with a Nikon Ti2E wide-field microscope.

*Figure generation* – Data plotting and statistical analyses were conducted using the Python SciPy, Matplotlib, and Seaborn packages. All figures were created in Inkscape v1.1.

## Acknowledgments

We thank Melanie Cobb for nomenclature suggestions; Josh Beck and Ben Weaver for helpful comments on the manuscript; the UT Southwestern Electron Microscopy and UT Southwestern Proteomics core facilities for assistance with data collection and analysis. M.L.R. acknowledges funding from the Welch Foundation (I-2075-20210327), National Science Foundation (MCB1553334), and NIH (AI150715). X.H. was funded, in part, by Cancer Prevention and Research Institute of Texas Training Grant RP160157. S.A.H. was funded, in part, by an NSF GRFP.

**Conflict of Interests**

The authors declare that they have no conflict of interest.

## Supplemental Material

**Supplemental Figure S1.**
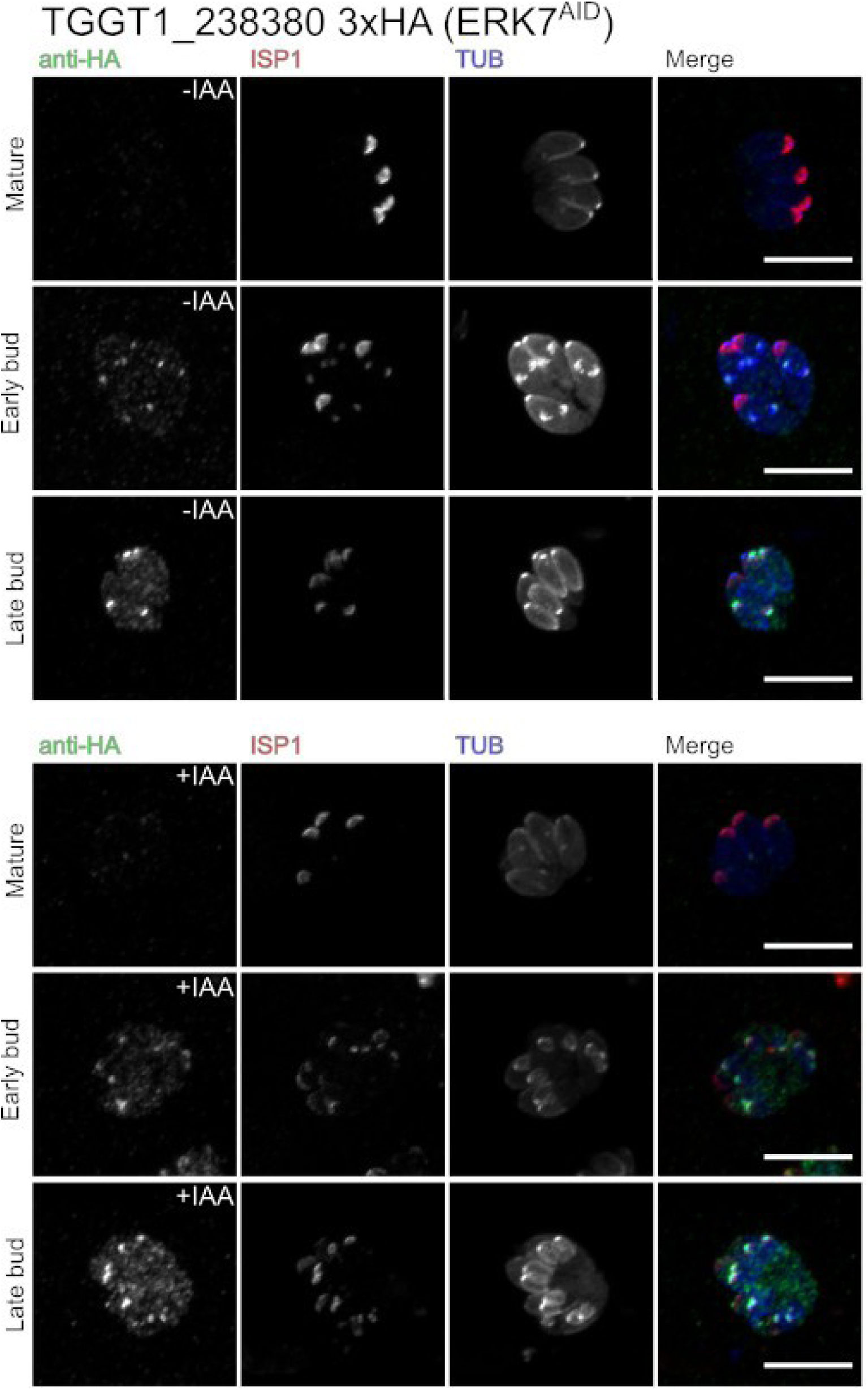
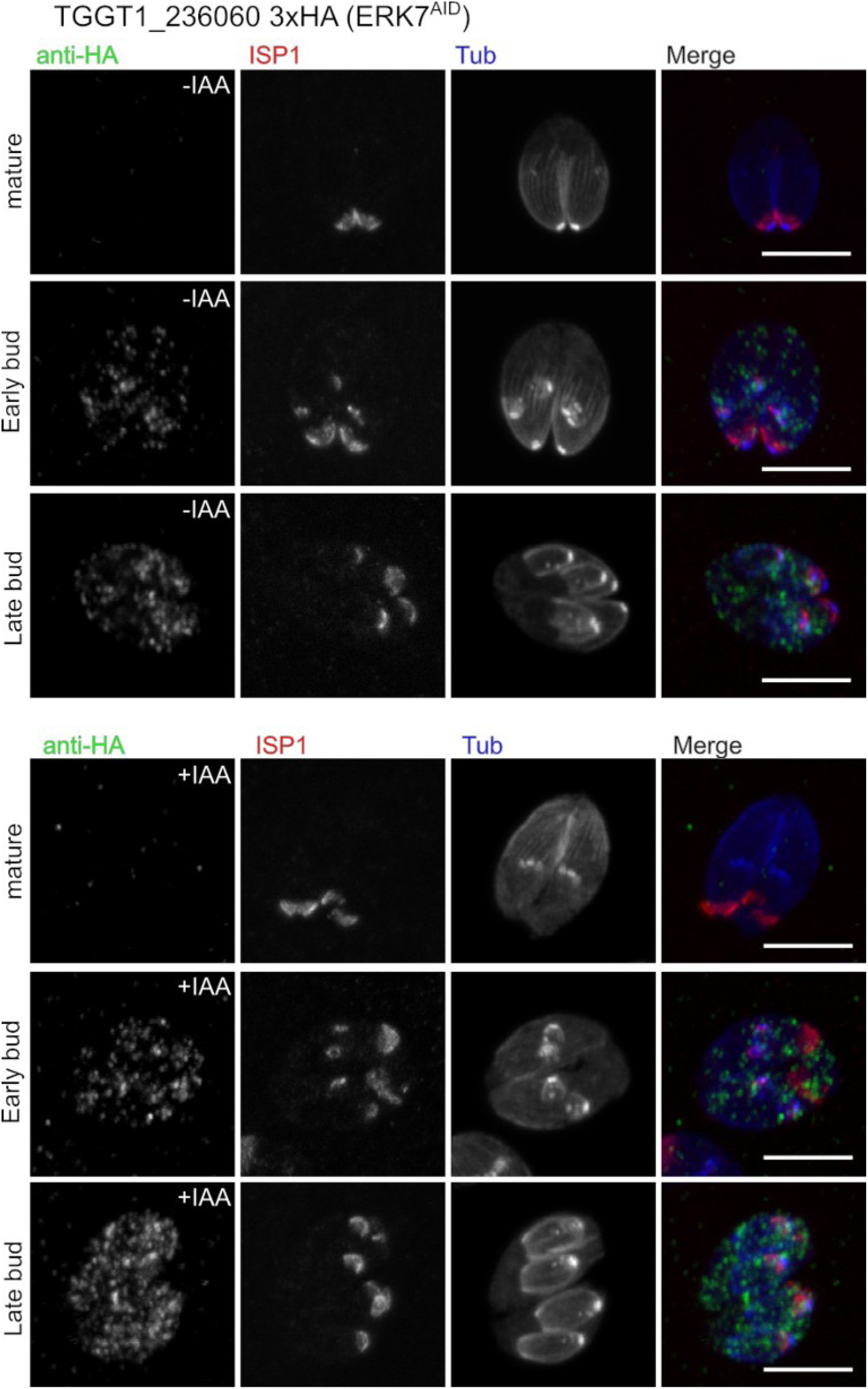
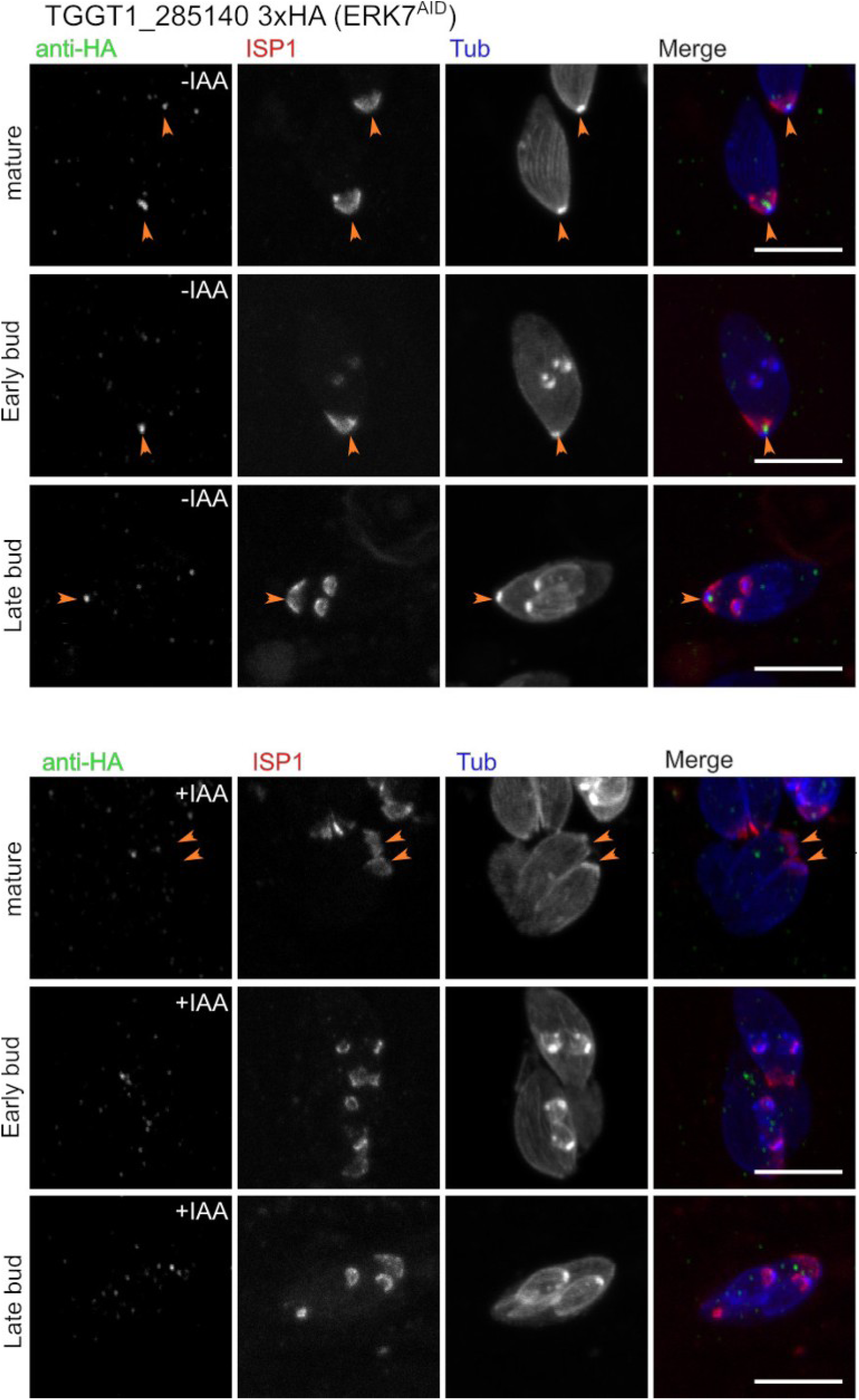
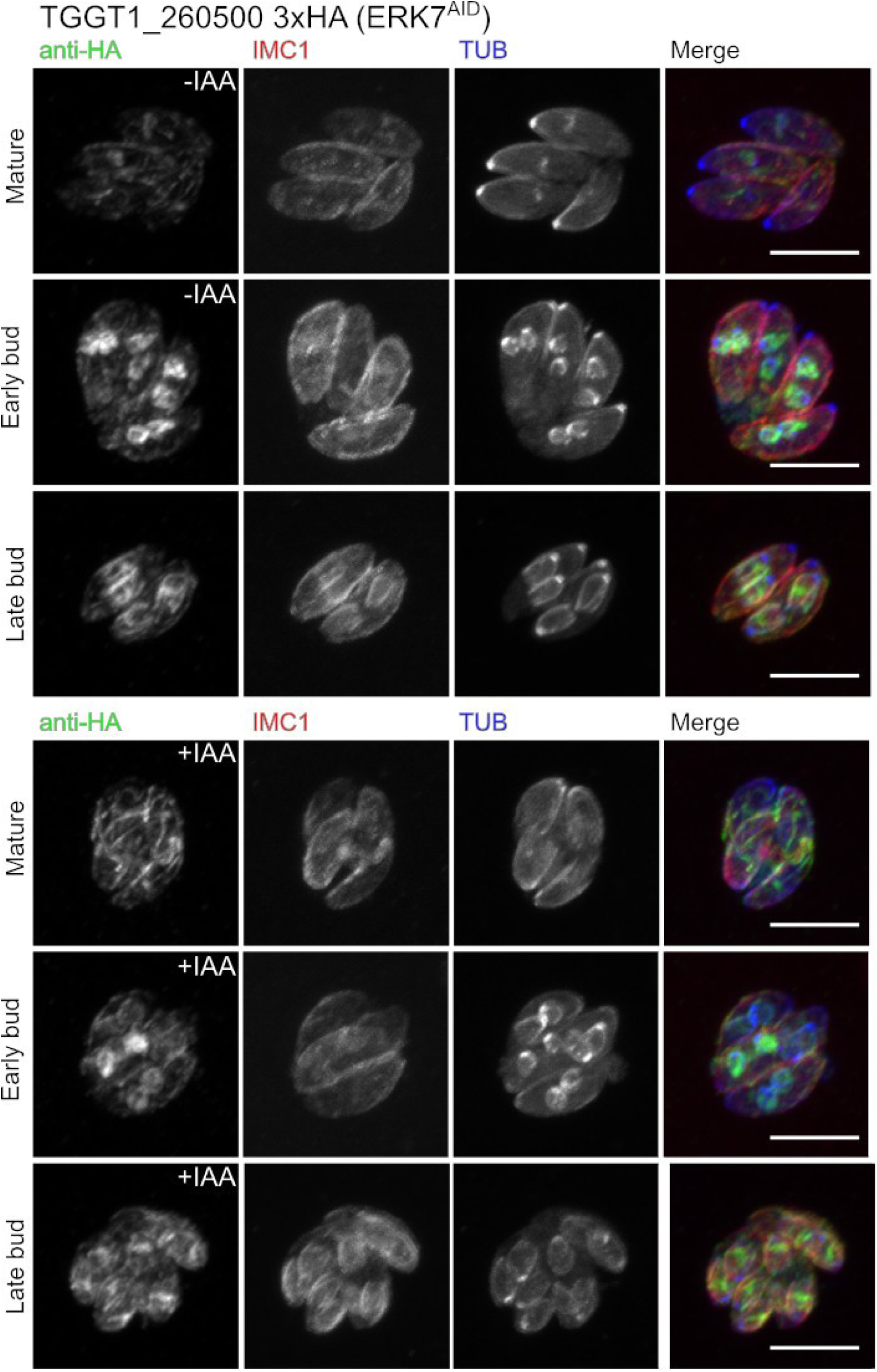
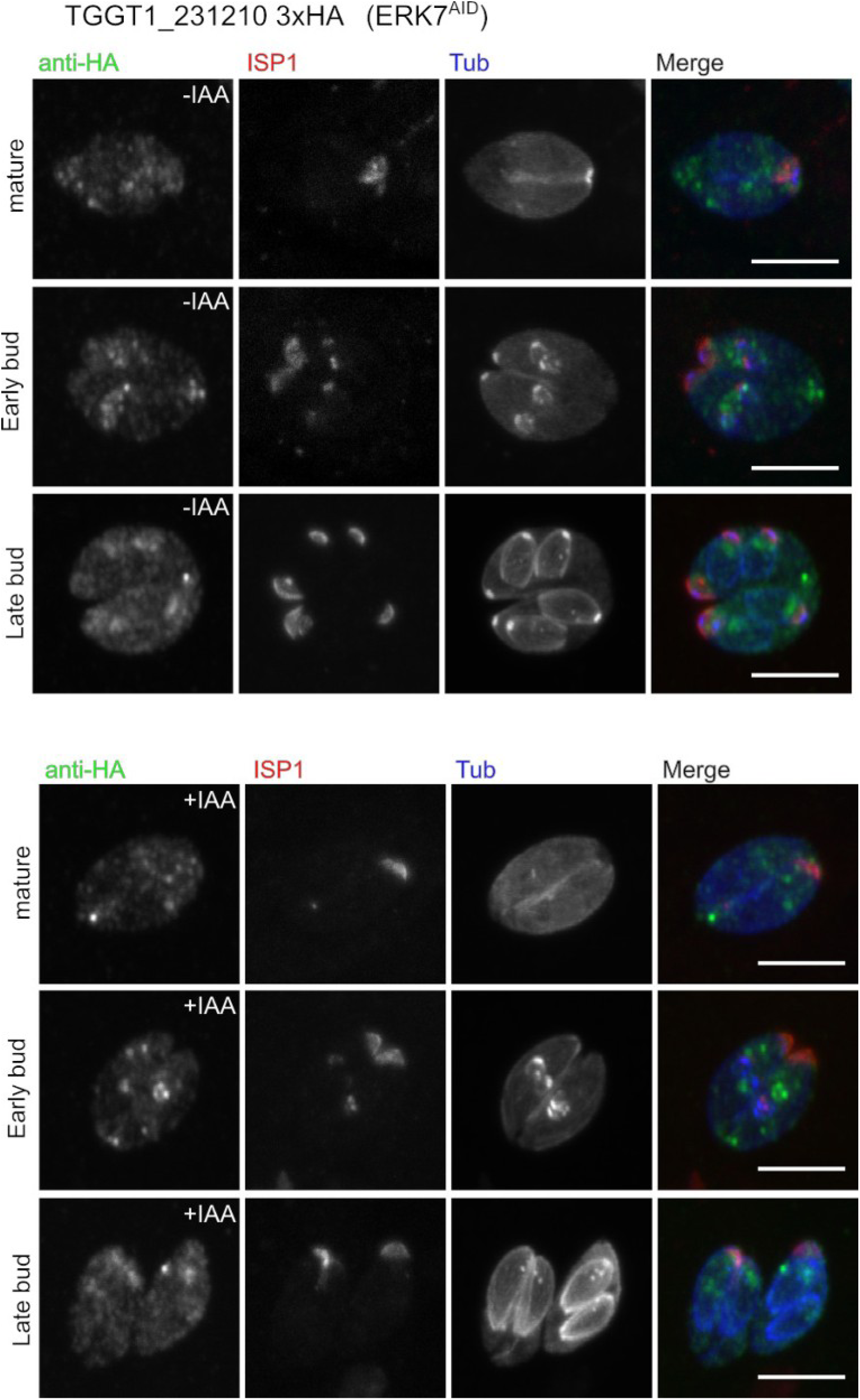
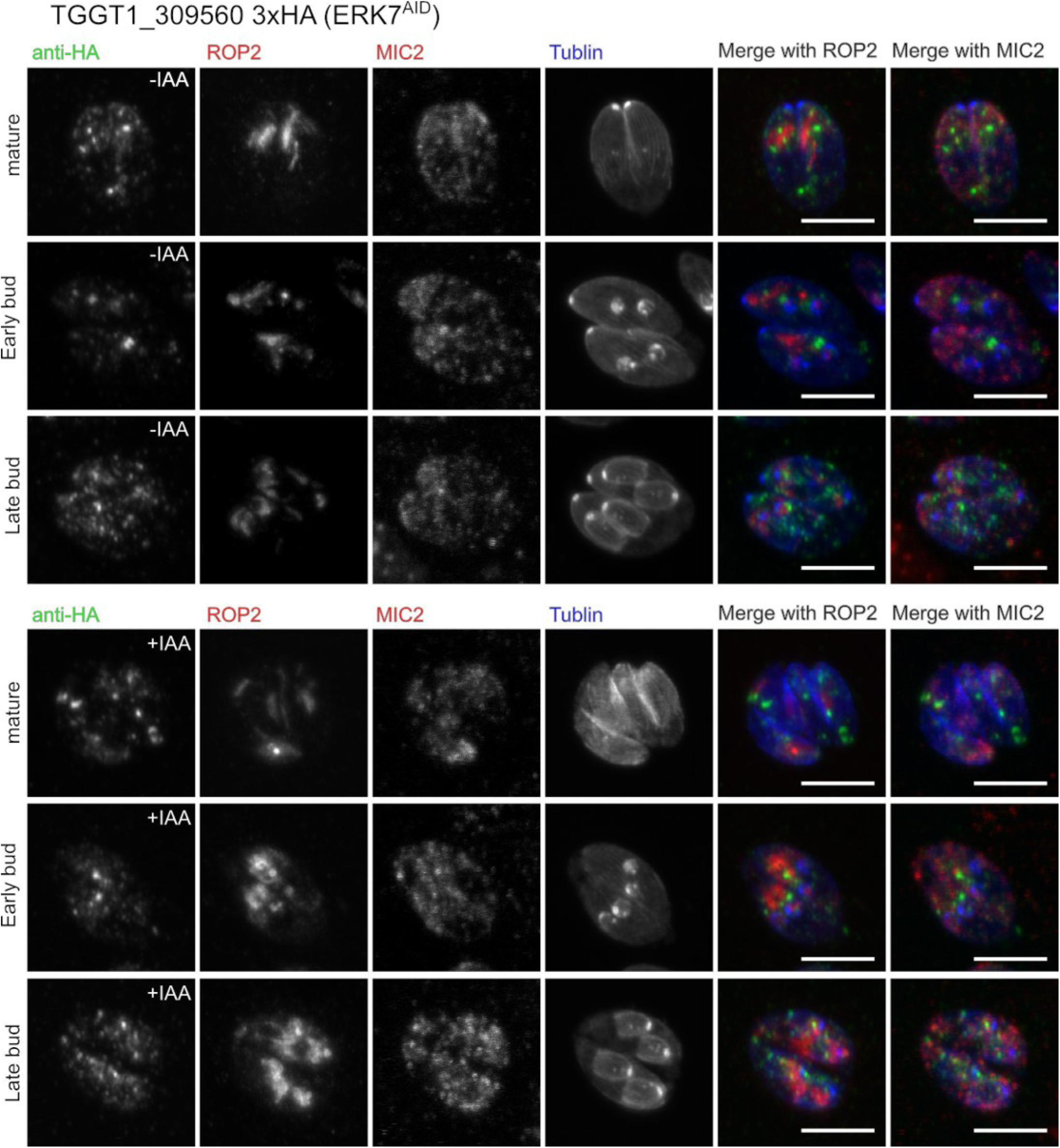
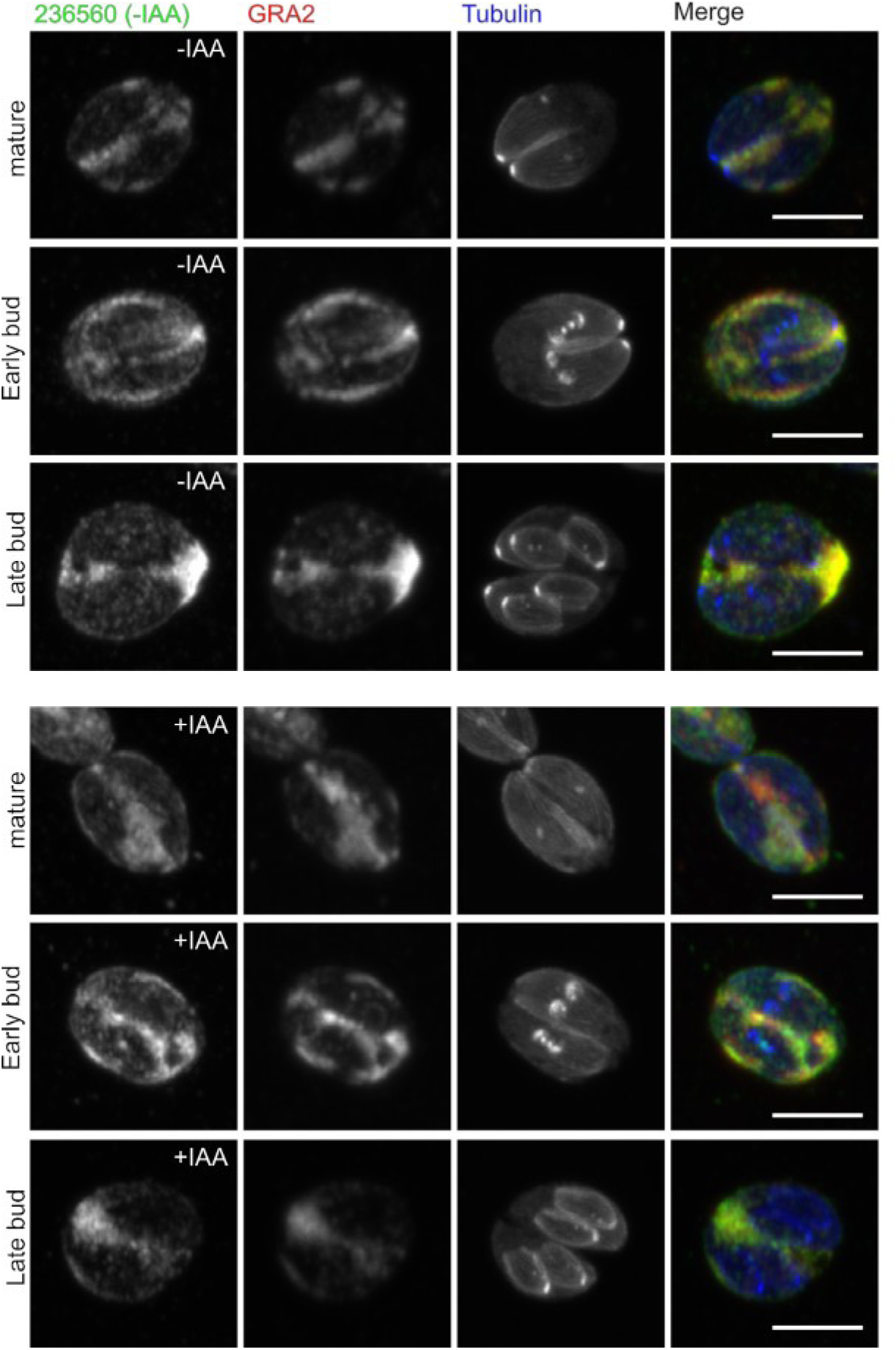
Candidates from the ERK7 interactome were examined by creating screens in which the gene of interest was expressed in-frame with a C-terminal 3xHA tag in the background of the ERK7^AID^ strain. Parasites were grown in ±IAA for 24 h before fixation. Samples were stained with antibodies recognizing the HA-tag and the ndicated counter-stain (usually ISP1, a marker for the apical cap). All images are representative of a minimum of n=20 images and are maximum intensity Z-projections of confocal stacks. **Supplemental Figure S1G:** While TGGT1_236560 was a hit in both our Yeast-two-hybrid and BioID datasets, C-terminal tagging led to vacuole localization, which is inconsistent with an interaction with cytosolic ERK7. TGGT1_236560 also has no predicted signal peptide. Note that the predicted protein length s ~250 kDa, and the gene has no predicted introns, which is unusual for a gene of this size. It is therefore possible that the gene model is incorrect and represents a combination of a cytosolic protein (at the 5’ end of the gene) and a secreted protein (at the 3’ end).

**Supplemental Figure S2:**
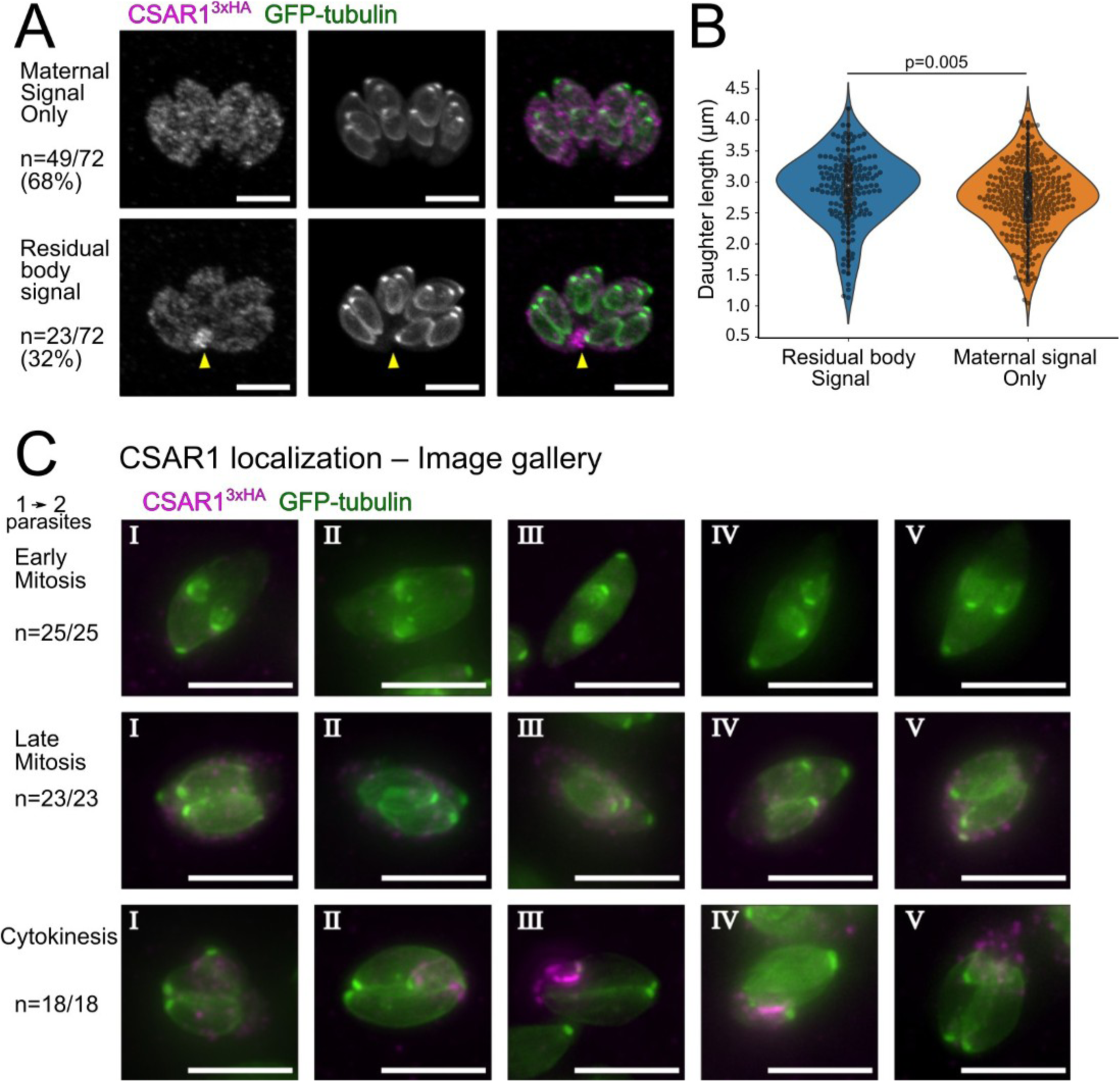
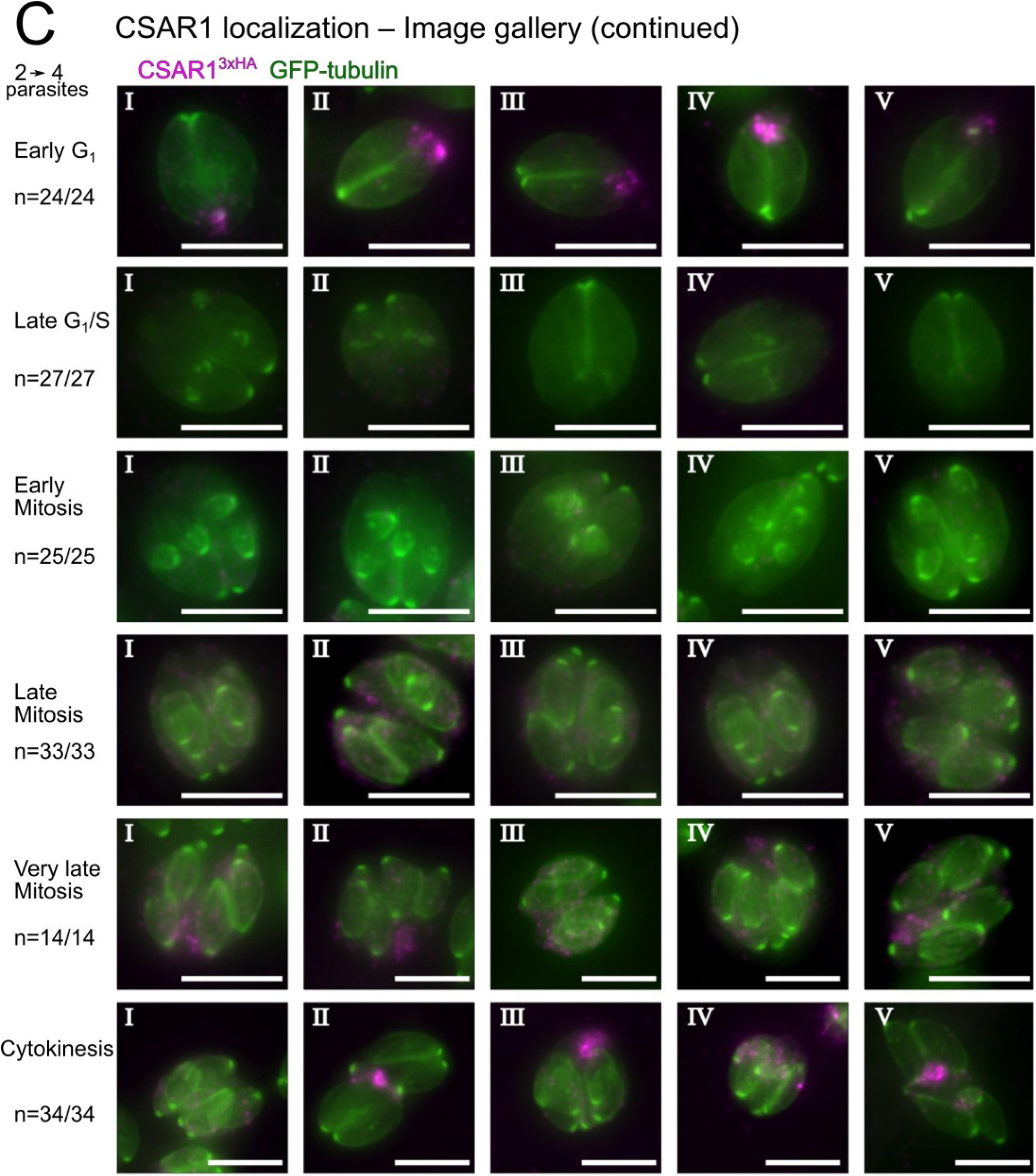
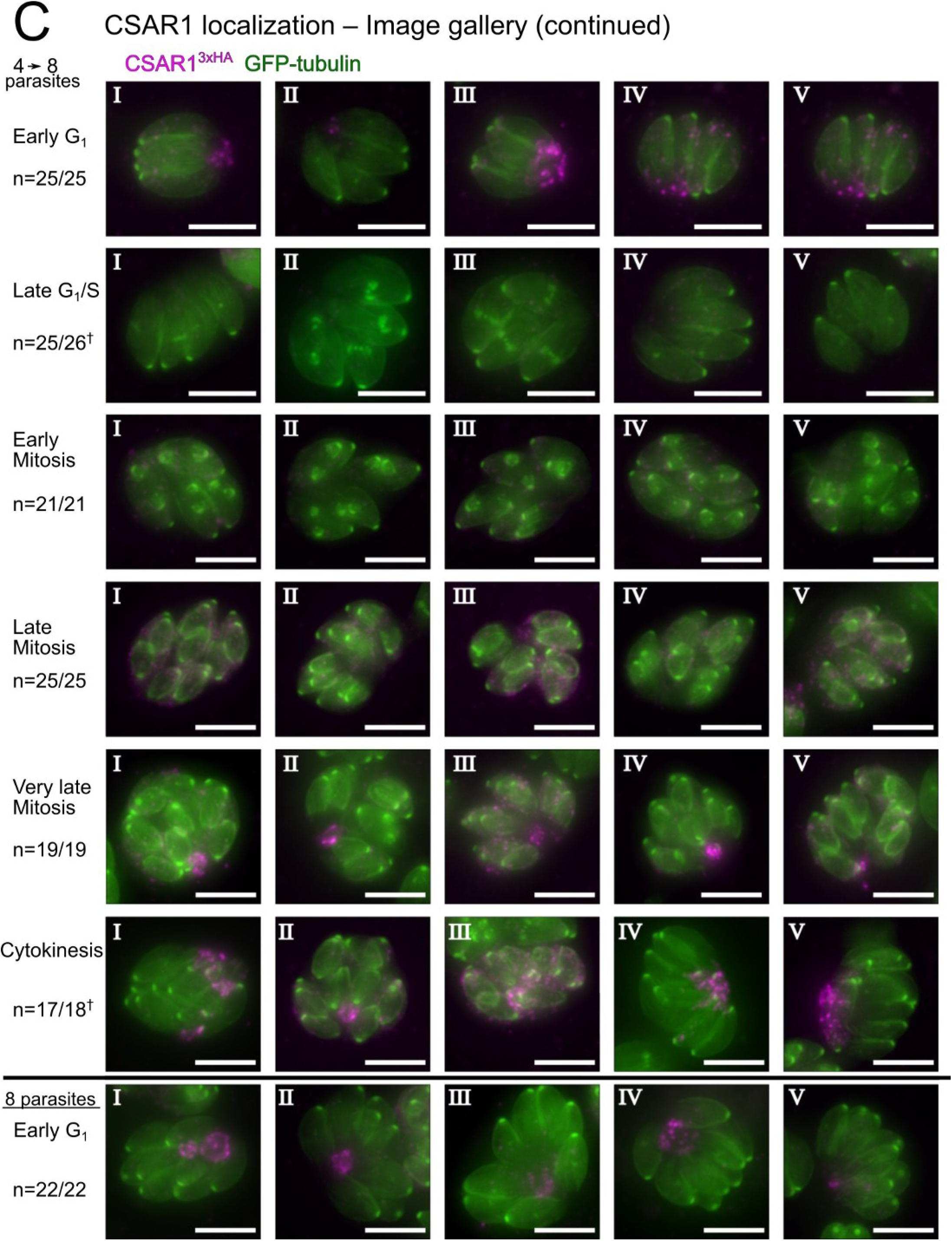
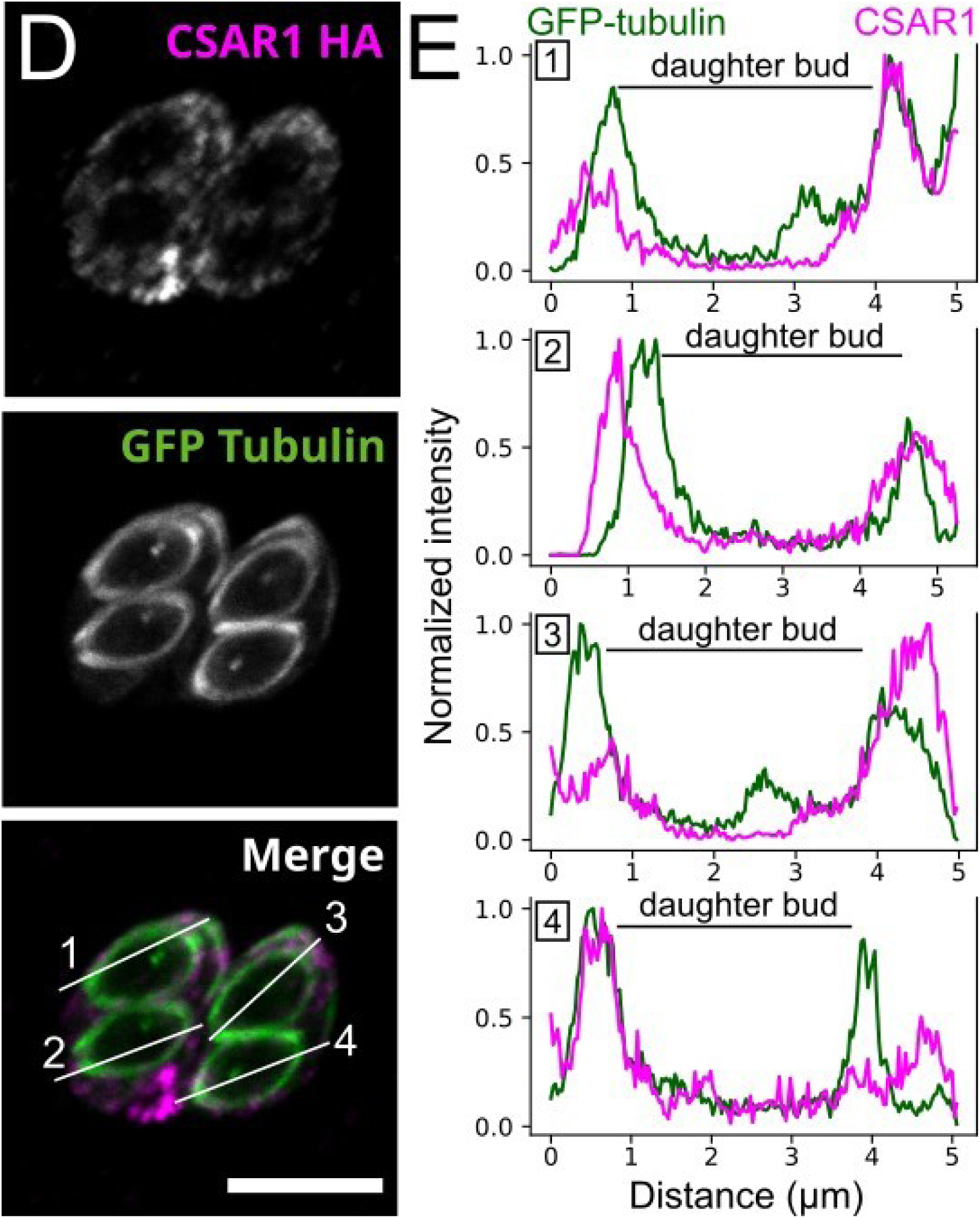
CSAR1 exhibits dynamic localization through the *Toxoplasma* cell cycle. (A) Late in mitosis (monitored by GFP-tubulin signal; green), in 68% of vacuoles examined, CSAR1^3xHA^ signal (magenta) is bright and confined to the maternal cytosol (see Supplemental Figure S3, below). In 32% of these vacuoles, CSAR1^3xHA^ has begun to concentrate at the residual body (lower panel; yellow arrowhead). As the vacuoles with CSAR1-residual body signal tend to contain larger daughter buds on average (B), we assume this relocalization of CSAR1 is occurring in parasites that have progressed further n mitosis, and which we therefore refer to in (C) as “Very late mitosis”. (C) Gallery of representative images of CSAR1^3xHA^ localization during cell cycle in parasites untreated with IAA. During the first replication cycle (1→2), there is no residual body present until cytokinesis. At this point CSAR1 signal concentrates at the site of the newly formed residual body, and co-localizes with the discarded maternal cytoskeleton. Localization was also tracked during the 2^nd^ and 3^rd^ divisions in images on the following 2 pages. Some groups contained parasites with morphologies consistent with dead or dying parasites, and these “n” are ndicated with (†). (D) 0.5 μm slice through a budding CSAR1^3xHA^ parasites from a confocal stack. CSAR1^3xHA^ (magenta), while building up in the maternal cytosol, appears excluded from the cytosol of daughter buds, which are outlined by GFP-tubulin (green). Scale bar is 5 μm. (E) A line scan at the indicated position in (D) highlights the lack of CSAR1^3xHA^ signal within a daughter bud. **Supplemental Figure S2**: **CSAR1 is excluded from daughter buds**. (D) 0.5 μm slice through a budding CSAR1^3xHA^ parasites from a confocal stack. CSAR1^3xHA^ (magenta), while building up in the maternal cytosol, appears excluded from the cytosol of daughter buds, which are outlined by GFP-tubulin (green). Scale bar s 5 μm. (E) A line scan at the indicated position in (D) highlights the lack of CSAR1^3xHA^ signal within a daughter bud.

**Supplemental Figure S3:**
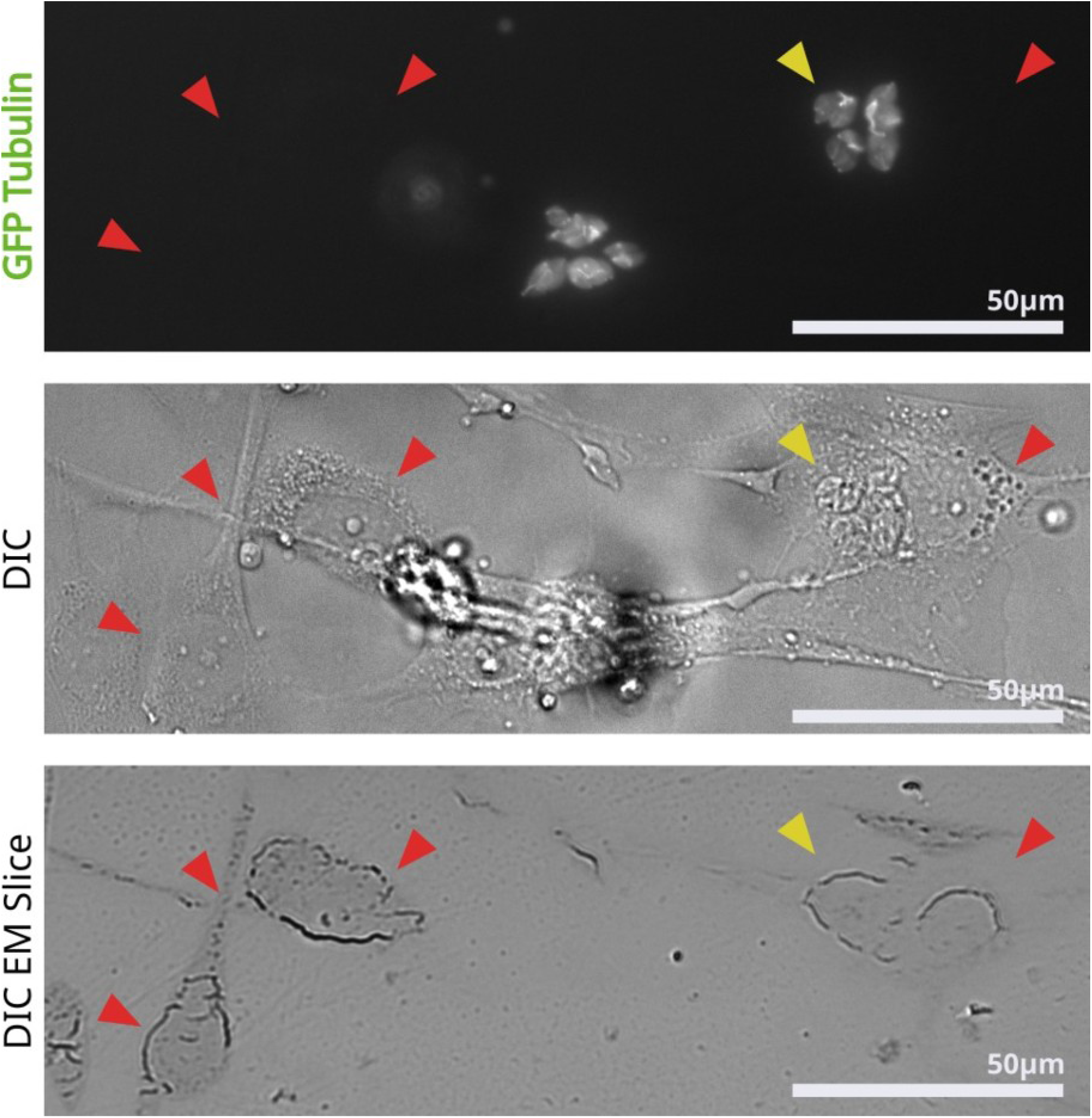
To generate samples for transmission electron microscopic analysis of retained cytoskeleton in *Δcsar1* vacuoles (see Figure 5), infected cells were first imaged by fluorescence (GFP-tubulin) and DIC to identify regions of interest. Multiple slices were then mounted on EM grids. Shown is an additional slice in this region mounted on a slide for visualization by DIC. Red arrowheads indicate landmark features. Yellow arrowhead indicates the cell infected with 4 vacuoles of *Δcsar1* parasites that was imaged and shown in Figure 5.

**Supplemental Figure S4.**
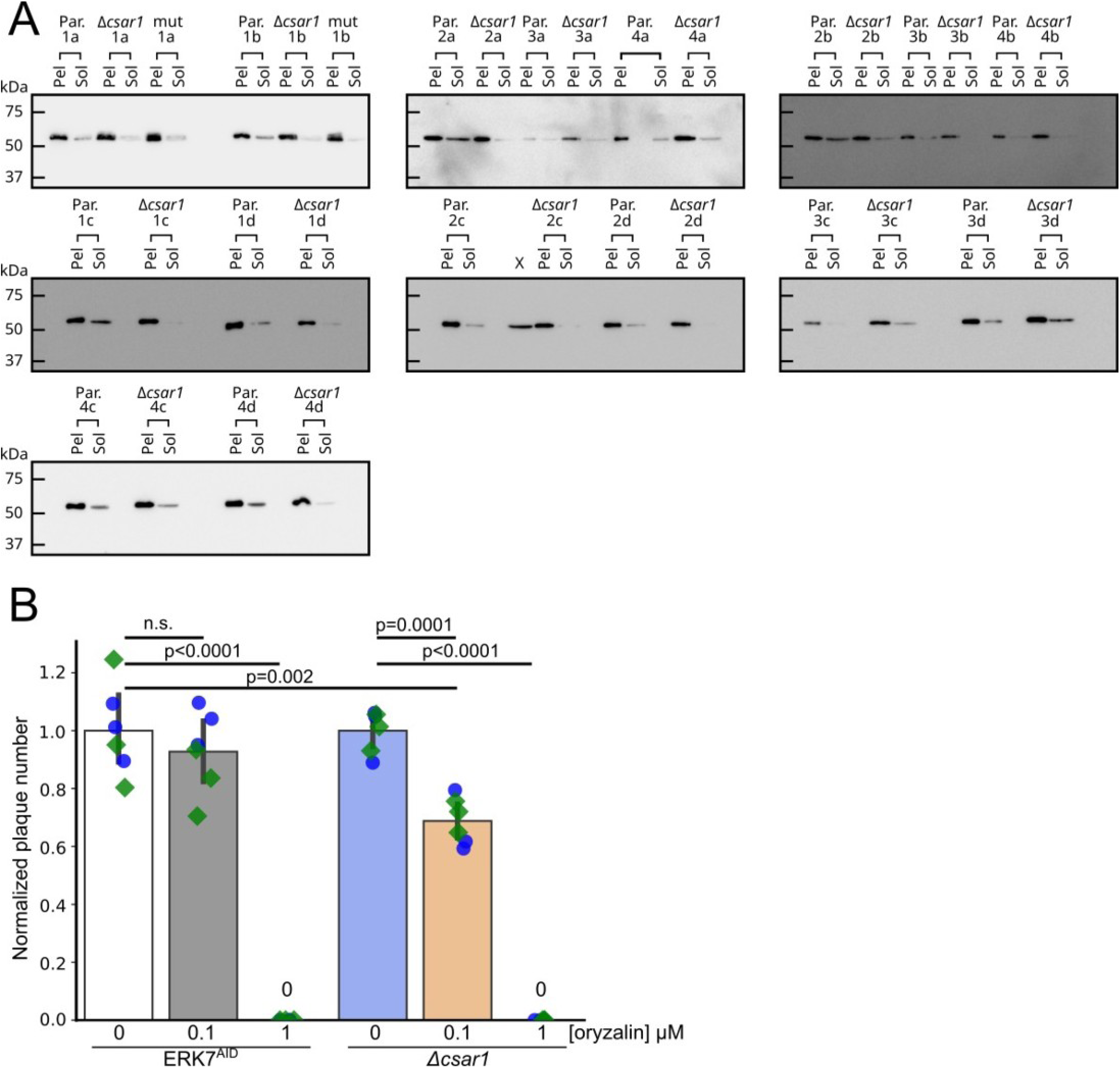
Loss of CSAR1 reduces the amount of soluble tubulin available for polymerization (A) Western blots quantified in Figure 6A. *Toxoplasma* polymerized tubulin structures are stable in detergent such as Triton-X-100, and can be separated from unpolymerized tubulin by centrifugation (see Methods). Equal volumes of soluble versus insoluble, assembled cytoskeleton from 4 biological replicates (1-4) with 4 technical replicates each (A-D) of either Parental (Par.) or *Δcsar1* parasites were separated by SDS-PAGE and quantified by western blot against *Toxoplasma* β-Tub. (B) Comparison the sensitivity of parental (ERK7^AID^) and *Δcsar1* parasites to the tubulin-polymerization inhibitor orazylin. Parasites were grown for 10 days in the indicated concentrations of oryzalin and the numbers of resulting plaques were normalized to the vehicle control. Significance was calculated by 1-way ANOVA followed by Tukey’s multiple comparison test (n.s., not significant).

**Supplemental Figure S5.**
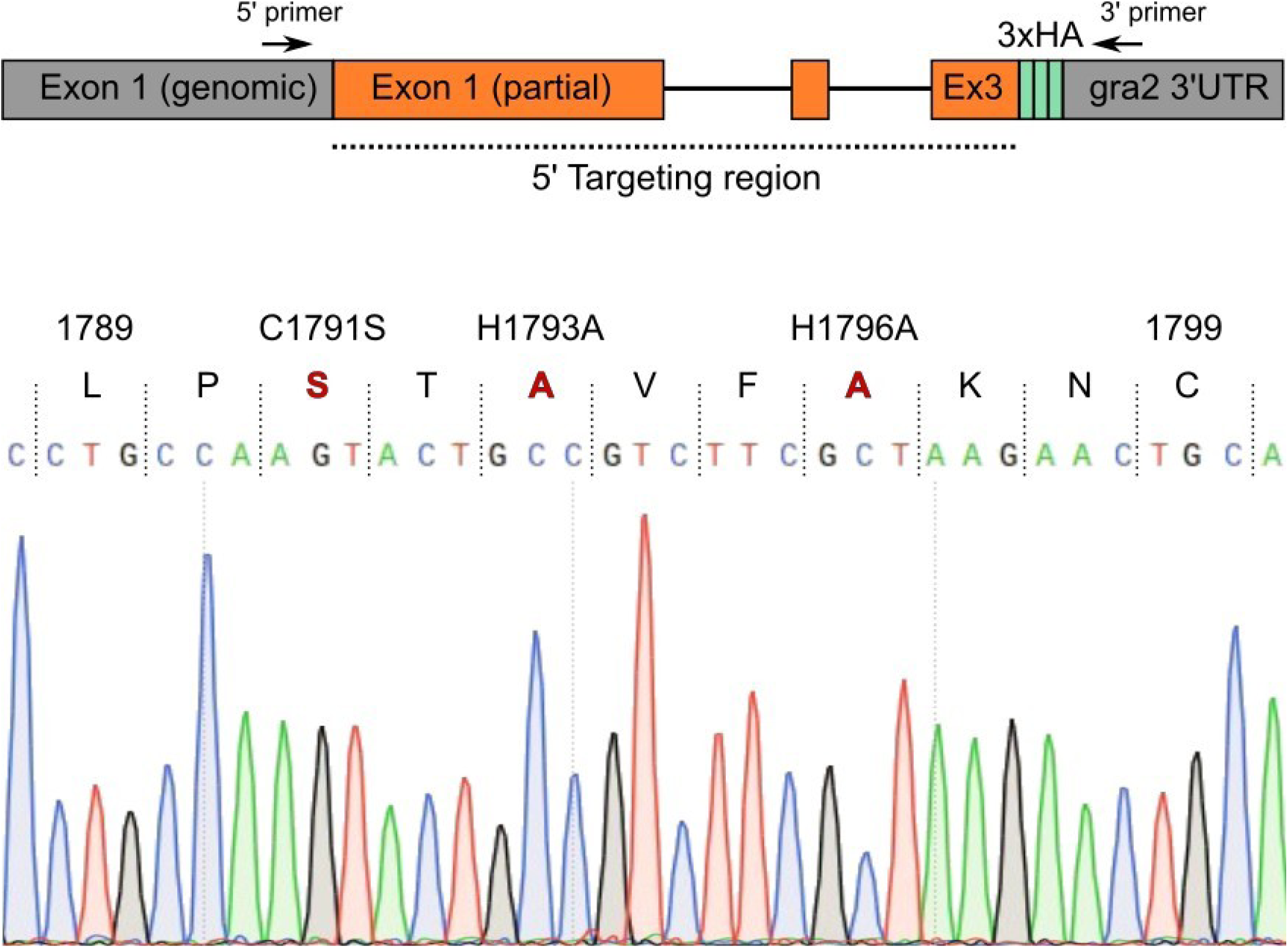
To validate the CSAR1 RING mutant, we amplified the 5’ end of the gene from the genomic DNA, using primers that annealed in the indicated regions. Note that the 5’ primer anneals in the csar1 locus outside of the targeting region of the construct used, to ensure that the amplicon results from the endogenous locus. 5’ primer (5’-ACATCGGCGGAGGAAGAGAG-3’), 3’ primer (5’-GTAGACTTCTCCCTTCAGCGGC-3’). The resulting amplicon was purified and then sequenced with the same primers. Sequencing confirmed correct integration into the endogenous locus and wild-type sequence outside of the area targeted for mutagenesis. A region of the resulting chromatogram is shown with the three mutated residues shown in red.

## References

1. M.-J. Gubbels, et al., Fussing About Fission: Defining Variety Among Mainstream and Exotic Apicomplexan Cell Division Modes. Front. Cell. Infect. Microbiol. 10, 269 (2020).

2. M.-J. Gubbels, I. Coppens, K. Zarringhalam, M. T. Duraisingh, K. Engelberg, The Modular Circuitry of Apicomplexan Cell Division Plasticity. Front. Cell. Infect. Microbiol. 11, 670049 (2021).

3. M. E. Francia, B. Striepen, Cell division in apicomplexan parasites. Nat. Rev. Microbiol. 12, 125–136 (2014).

4. M. Nishi, K. Hu, J. M. Murray, D. S. Roos, Organellar dynamics during the cell cycle of Toxoplasma gondii. J. Cell Sci. 121, 1559–1568 (2008).

5. L. Pelletier, et al., Golgi biogenesis in Toxoplasma gondii. Nature 418, 548–552 (2002).

6. D. T. Riglar, et al., Spatial association with PTEX complexes defines regions for effector export into Plasmodium falciparum-infected erythrocytes. Nat. Commun. 4, 1415 (2013).

7. J. Q. Tran, et al., RNG1 is a late marker of the apical polar ring in Toxoplasma gondii. Cytoskelet. Hoboken NJ 67, 586–598 (2010).

8. B. R. Anderson-White, et al., A family of intermediate filament-like proteins is sequentially assembled into the cytoskeleton of Toxoplasma gondii. Cell. Microbiol. 13, 18–31 (2011).

9. B. Anderson-White, et al., Cytoskeleton assembly in Toxoplasma gondii cell division. Int. Rev. Cell Mol. Biol. 298, 1–31 (2012).

10. M. Goldman, R. K. Carver, A. J. Sulzer, Reproduction of Toxoplasma gondii by internal budding. J. Parasitol. 44, 161–171 (1958).

11. M. A. Gavin, T. Wanko, L. Jacobs, Electron microscope studies of reproducing and interkinetic Toxoplasma. J. Protozool. 9, 222–234 (1962).

12. M. Attias, K. Miranda, W. De Souza, Development and fate of the residual body of Toxoplasma gondii. Exp. Parasitol. 196, 1–11 (2019).

13. J. Periz, et al., Toxoplasma gondii F-actin forms an extensive filamentous network required for material exchange and parasite maturation. eLife 6 (2017).

14. N. Tosetti, N. Dos Santos Pacheco, D. Soldati-Favre, D. Jacot, Three F-actin assembly centers regulate organelle inheritance, cell-cell communication and motility in Toxoplasma gondii. eLife 8 (2019).

15. E. S. Suvorova, M. Francia, B. Striepen, M. W. White, A novel bipartite centrosome coordinates the apicomplexan cell cycle. PLoS Biol. 13, e1002093 (2015).

16. C.-T. Chen, M.-J. Gubbels, TgCep250 is dynamically processed through the division cycle and is essential for structural integrity of the Toxoplasma centrosome. Mol. Biol. Cell 30, 1160–1169 (2019).

17. K. Hu, et al., Daughter cell assembly in the protozoan parasite Toxoplasma gondii. Mol. Biol. Cell 13, 593–606 (2002).

18. B. Striepen, et al., The plastid of Toxoplasma gondii is divided by association with the centrosomes. J. Cell Biol. 151, 1423–1434 (2000).

19. C. A. Alvarez, E. S. Suvorova, Checkpoints of apicomplexan cell division identified in Toxoplasma gondii. PLoS Pathog. 13, e1006483 (2017).

20. X. Hu, W. J. O’Shaughnessy, T. G. Beraki, M. L. Reese, Loss of the Conserved Alveolate Kinase MAPK2 Decouples Toxoplasma Cell Growth from Cell Division. mBio 11 (2020).

21. C.-T. Chen, M.-J. Gubbels, The Toxoplasma gondii centrosome is the platform for internal daughter budding as revealed by a Nek1 kinase mutant. J. Cell Sci. 126, 3344–3355 (2013).

22. L. Koreny, et al., Molecular characterization of the conoid complex in Toxoplasma reveals its conservation in all apicomplexans, including Plasmodium species. PLoS Biol. 19, e3001081 (2021).

23. E. Bertiaux, et al., Expansion microscopy provides new insights into the cytoskeleton of malaria parasites including the conservation of a conoid. PLoS Biol. 19, e3001020 (2021).

24. Z. Füssy, P. Masařová, J. Kručinská, H. J. Esson, M. Oborník, Budding of the Alveolate Alga Vitrella brassicaformis Resembles Sexual and Asexual Processes in Apicomplexan Parasites. Protist 168, 80–91 (2017).

25. N. Okamoto, P. J. Keeling, The 3D structure of the apical complex and association with the flagellar apparatus revealed by serial TEM tomography in Psammosa pacifica, a distant relative of the Apicomplexa. PloS One 9, e84653 (2014).

26. K. Hu, D. S. Roos, J. M. Murray, A novel polymer of tubulin forms the conoid of Toxoplasma gondii. J. Cell Biol. 156, 1039–1050 (2002).

27. J. C. de Leon, et al., A SAS-6-like protein suggests that the Toxoplasma conoid complex evolved from flagellar components. Eukaryot. Cell 12, 1009–1019 (2013).

28. M. E. Francia, et al., Cell division in Apicomplexan parasites is organized by a homolog of the striated rootlet fiber of algal flagella. PLoS Biol. 10, e1001444 (2012).

29. M. E. Francia, J.-F. Dubremetz, N. S. Morrissette, Basal body structure and composition in the apicomplexans Toxoplasma and Plasmodium. Cilia 5, 3 (2015).

30. G. Lentini, D. J. Dubois, B. Maco, D. Soldati-Favre, K. Frénal, The roles of Centrin 2 and Dynein Light Chain 8a in apical secretory organelles discharge of Toxoplasma gondii. Traffic Cph. Den. 20, 583–600 (2019).

31. M. F. Lévêque, L. Berry, S. Besteiro, An evolutionarily conserved SSNA1/DIP13 homologue is a component of both basal and apical complexes of Toxoplasma gondii. Sci. Rep. 6, 27809 (2016).

32. M. G. Del Carmen, M. Mondragón, S. González, R. Mondragón, Induction and regulation of conoid extrusion in Toxoplasma gondii. Cell. Microbiol. 11, 967–982 (2009).

33. V. B. Carruthers, L. D. Sibley, Mobilization of intracellular calcium stimulates microneme discharge in Toxoplasma gondii. Mol. Microbiol. 31, 421–428 (1999).

34. A. Graindorge, et al., The Conoid Associated Motor MyoH Is Indispensable for Toxoplasma gondii Entry and Exit from Host Cells. PLoS Pathog. 12, e1005388 (2016).

35. M. K. Shaw, H. L. Compton, D. S. Roos, L. G. Tilney, Microtubules, but not actin filaments, drive daughter cell budding and cell division in Toxoplasma gondii. J. Cell Sci. 113 (Pt 7), 1241–1254 (2000).

36. B. A. Nichols, M. L. Chiappino, Cytoskeleton of Toxoplasma gondii. J. Protozool. 34, 217–226 (1987).

37. W. J. O’Shaughnessy, X. Hu, T. Beraki, M. McDougal, M. L. Reese, Loss of a conserved MAPK causes catastrophic failure in assembly of a specialized cilium-like structure in Toxoplasma gondii. Mol. Biol. Cell 31, 881–888 (2020).

38. K. Nishimura, T. Fukagawa, H. Takisawa, T. Kakimoto, M. Kanemaki, An auxin-based degron system for the rapid depletion of proteins in nonplant cells. Nat. Methods 6, 917–22 (2009).

39. P. S. Back, et al., Ancient MAPK ERK7 is regulated by an unusual inhibitory scaffold required for Toxoplasma apical complex biogenesis. Proc. Natl. Acad. Sci. U. S. A. 117, 12164–12173 (2020).

40. N. Tosetti, et al., Essential function of the alveolin network in the subpellicular microtubules and conoid assembly in Toxoplasma gondii. eLife 9 (2020).

41. D. Sang, et al., Ancestral reconstruction reveals mechanisms of ERK regulatory evolution. eLife 8 (2019).

42. K. J. Roux, D. I. Kim, B. Burke, BioID: a screen for protein-protein interactions. Curr. Protoc. Protein Sci. 74, Unit 19.23. (2013).

43. S. Fields, O. Song, A novel genetic system to detect protein-protein interactions. Nature 340, 245–246 (1989).

44. K. J. Roux, D. I. Kim, M. Raida, B. Burke, A promiscuous biotin ligase fusion protein identifies proximal and interacting proteins in mammalian cells. J. Cell Biol. 196, 801–10 (2012).

45. D. I. Kim, et al., Probing nuclear pore complex architecture with proximity-dependent biotinylation. Proc. Natl. Acad. Sci. U. S. A. 111, E2453–2461 (2014).

46. D. I. Kim, et al., An improved smaller biotin ligase for BioID proximity labeling. Mol. Biol. Cell 27, 1188–1196 (2016).

47. M. S. Behnke, et al., Coordinated progression through two subtranscriptomes underlies the tachyzoite cycle of Toxoplasma gondii. PLoS One 5, e12354 (2010).

48. K. Hu, et al., Cytoskeletal components of an invasion machine--the apical complex of Toxoplasma gondii. PLoS Pathog. 2, e13 (2006).

49. P. S. Back, et al., Multivalent Interactions Drive the Toxoplasma AC9:AC10:ERK7 Complex To Concentrate ERK7 in the Apical Cap. mBio, e0286421 (2022).

50. N. S. Morrissette, L. D. Sibley, Disruption of microtubules uncouples budding and nuclear division in Toxoplasma gondii. J. Cell Sci. 115, 1017–1025 (2002).

51. A. Akhmanova, M. O. Steinmetz, Control of microtubule organization and dynamics: two ends in the limelight. Nat. Rev. Mol. Cell Biol. 16, 711–726 (2015).

52. K. Hu, D. S. Roos, S. O. Angel, J. M. Murray, Variability and heritability of cell division pathways in Toxoplasma gondii. J. Cell Sci. 117, 5697–5705 (2004).

53. R. J. Deshaies, C. A. P. Joazeiro, RING domain E3 ubiquitin ligases. Annu. Rev. Biochem. 78, 399–434 (2009).

54. K. Miyatake, M. Kusakabe, C. Takahashi, E. Nishida, ERK7 regulates ciliogenesis by phosphorylating the actin regulator CapZIP in cooperation with Dishevelled. Nat. Commun. 6, 6666 (2015).

55. A. Kazatskaya, et al., Primary Cilium Formation and Ciliary Protein Trafficking Is Regulated by the Atypical MAP Kinase MAPK15 in Caenorhabditis elegans and Human Cells. Genetics 207, 1423–1440 (2017).

56. Y. Wei, Z. Li, Distinct roles of a mitogen-activated protein kinase in cytokinesis between different life cycle forms of Trypanosoma brucei. Eukaryot. Cell 13, 110–118 (2014).

57. M. Zacharogianni, et al., ERK7 is a negative regulator of protein secretion in response to amino-acid starvation by modulating Sec16 membrane association. EMBO J. 30, 3684–3700 (2011).

58. J. A. Brzostowski, et al., Phosphorylation of chemoattractant receptors regulates chemotaxis, actin reorganization and signal relay. J. Cell Sci. 126, 4614–4626 (2013).

59. J. Chia, K. M. Tham, D. J. Gill, E. A. Bard-Chapeau, F. A. Bard, ERK8 is a negative regulator of O-GalNAc glycosylation and cell migration. eLife 3, e01828 (2014).

60. D. P. Bermingham, et al., The Atypical MAP Kinase SWIP-13/ERK8 Regulates Dopamine Transporters through a Rho-Dependent Mechanism. J. Neurosci. Off. J. Soc. Neurosci. 37, 9288–9304 (2017).

61. D. Colecchia, et al., MAPK15/ERK8 stimulates autophagy by interacting with LC3 and GABARAP proteins. Autophagy 8, 1724–1740 (2012).

62. D. T. Ouologuem, D. S. Roos, Dynamics of the Toxoplasma gondii inner membrane complex. J. Cell Sci. 127, 3320–3330 (2014).

63. R. Dubey, et al., Differential Roles for Inner Membrane Complex Proteins across Toxoplasma gondii and Sarcocystis neurona Development. mSphere 2, e00409–17 (2017).

64. J. R. Beck, et al., A novel family of Toxoplasma IMC proteins displays a hierarchical organization and functions in coordinating parasite division. PLoS Pathog. 6, e1001094 (2010).

65. A. Hunt, et al., Differential requirements for cyclase-associated protein (CAP) in actin-dependent processes of Toxoplasma gondii. eLife 8 (2019).

66. M. Huynh, V. B. Carruthers, Tagging of endogenous genes in a Toxoplasma gondii strain lacking Ku80. Eukaryot. Cell 8, 530–9 (2009).

67. J. Schindelin, et al., Fiji: an open-source platform for biological-image analysis. Nat. Methods 9, 676–682 (2012).

68. E. Nagayasu, Y.-C. Hwang, J. Liu, J. M. Murray, K. Hu, Loss of a doublecortin (DCX)-domain protein causes structural defects in a tubulin-based organelle of Toxoplasma gondii and impairs host-cell invasion. Mol. Biol. Cell 28, 411–428 (2017).

69. A. L. Chen, et al., Novel insights into the composition and function of the Toxoplasma IMC sutures. Cell. Microbiol. 19 (2017).

70. A. Lorestani, et al., Targeted proteomic dissection of Toxoplasma cytoskeleton sub-compartments using MORN1. Cytoskeleton 69, 1069–1085 (2012).

71. C. G. Baptista, et al., Toxoplasma F-box protein 1 is required for daughter cell scaffold function during parasite replication. PLOS Pathog. 15, e1007946 (2019).

72. T. Mann, C. Beckers, Characterization of the subpellicular network, a filamentous membrane skeletal component in the parasite Toxoplasma gondii. Mol. Biochem. Parasitol. 115, 257–268 (2001).

73. M. J. Wichroski, J. A. Melton, C. G. Donahue, R. K. Tweten, G. E. Ward, Clostridium septicum alpha-toxin is active against the parasitic protozoan Toxoplasma gondii and targets members of the SAG family of glycosylphosphatidylinositol-anchored surface proteins. Infect. Immun. 70, 4353–4361 (2002).

74. S. B. Gould, W.-H. Tham, A. F. Cowman, G. I. McFadden, R. F. Waller, Alveolins, a new family of cortical proteins that define the protist infrakingdom Alveolata. Mol. Biol. Evol. 25, 1219–1230 (2008).

75. S. B. Gould, et al., Ciliate pellicular proteome identifies novel protein families with characteristic repeat motifs that are common to alveolates. Mol. Biol. Evol. 28, 1319–1331 (2011).

76. T. A. Smith, G. S. Lopez-Perez, A. L. Herneisen, E. Shortt, S. Lourido, Screening the Toxoplasma kinome with high-throughput tagging identifies a regulator of invasion and egress. Nat. Microbiol. 7, 868–881 (2022).

77. S. Long, B. Anthony, L. L. Drewry, L. D. Sibley, A conserved ankyrin repeat-containing protein regulates conoid stability, motility and cell invasion in Toxoplasma gondii. Nat. Commun. 8, 2236 (2017).

